# Personalized models of breast cancer desmoplasia reveal biomechanical determinants of drug penetration

**DOI:** 10.1101/2021.12.12.472296

**Authors:** Giovanni S. Offeddu, Elena Cambria, Sarah E. Shelton, Zhengpeng Wan, Kristina Haase, Luca Possenti, Huu Tuan Nguyen, Mark R. Gillrie, Dean Hickman, Charles G. Knutson, Roger D. Kamm

## Abstract

Breast cancer desmoplasia heterogeneity contributes to high disease mortality due to discrepancies in treatment efficacy between patients. Personalized *in vitro* breast cancer models can be used for high throughput testing and ranking of therapeutic strategies to normalize the aberrant microenvironment in a patient-specific manner. Here, tumoroids assembled from patient-derived cells cultured in microphysiological systems including perfusable microvasculature reproduce key aspects of stromal and vascular dysfunction. Increased hyaluronic acid and collagen deposition, loss of vascular glycocalyx and reduced perfusion, and elevated interstitial fluid pressure in the models result in impaired drug distribution to tumor cells. We demonstrate the application of these personalized models as tools to rank molecular therapies for the normalization of the tumoroid microenvironment and to discover new therapeutic targets such as IL8 and CD44, which may ultimately improve drug efficacy in breast cancer patients.

## 1. INTRODUCTION

Tissue desmoplasia is a key process underlying disease progression in breast cancer. The induced physical heterogeneity in the breast tissue presents a major obstacle to treatment^1,2^. Breast tumor cells (TCs) are largely responsible for desmoplastic remodeling of the tumor microenvironment, both directly through aberrant deposition of matrix proteins and indirectly through activation of stromal cells^3–5^. Increased ECM density, reduced vascular perfusion and barrier function, and reduced lymphatic drainage all contribute to increased interstitial fluid pressure (IFP) in the breast cancer microenvironment, resulting in reduced immune cell infiltration^6^ and impaired drug distribution from blood to TCs^7–9^. Therapeutic strategies to normalize the desmoplastic breast cancer microenvironment are currently in clinical trials^10^. However, the patient-to-patient heterogeneity of breast cancer, hence its response to different treatments, makes it critical to identify personalized strategies with the greatest therapeutic potential.

Patient-derived TCs formed into structures with increasing cellular complexity – single-cell type spheroids^11^, multi-cell type tumoroids^12,13^, and stem cell-derived organoids^14^ - have recently attracted attention as laboratory models to rapidly evaluate response to molecular therapies^15^. Incorporating various “-oid” structures into microphysiological models presents the opportunity to broaden their scope and capture key morphological and functional aspects of the tumor microenvironment^16–19^. An additional attractive feature of microfluidic-based models is the extremely fine control over mechanical and biochemical stimuli imparted on the cells within^20,21^. However, current models have so far been unsuccessful in recapitulating the complexity of the desmoplastic breast cancer microenvironment^22^, particularly the aberrant ECM and vasculature, which jointly determine drug distribution into the tumor. Harnessing the capabilities of microphysiological models to recapitulate the pathophysiological complexity of the aberrant TC microenvironment in breast cancer desmoplasia could provide new therapeutic strategies to improve clinical care for breast cancer patients.

We have recently demonstrated culture of cancer spheroids assembled from ovarian and lung TC lines in microphysiological models containing human microvascular networks (MVNs)^23^. These MVNs can be perfused with therapeutic molecules to assess their permeability across the vascular endothelium and resulting TC death. In this work, we expand on this methodology to culture tumoroids containing primary breast TCs or breast TC lines, evaluating their microenvironments to demonstrate differential desmoplastic stromal and vascular remodeling with resulting responsiveness to normalizing treatments. Herein, we show that these vascularized tumor models can both provide quantitative metrics to rank the efficacy of different therapeutic strategies, and empower the discovery of new molecular targets to normalize the desmoplastic breast cancer microenvironment.

## 2. RESULTS

### 2.1 Vascularized tumoroids on-chip reproduce variations in breast cancer desmoplasia

We first made use of breast TC lines representative of three main disease molecular sub-types^24^ to establish the model’s capabilities: MCF7 (estrogen receptor, ER, and progesterone receptor, PR, positive), SKBR3 (human epidermal growth factor receptor 2, HER2, positive) and MDA-MB-468 (“triple-negative”, epidermal growth factor receptor, EGFR, positive). Tumoroids approximately 500 µm in diameter were formed by co-culture of TCs with human fibroblasts (FBs) in non-adherent well plates for 4 days, which allowed the formation of stable cell aggregates even for TCs that would not aggregate in mono-culture (SKBR3 and MDA-MB-468). We employed our recently described^23^ methodology to co-culture the tumoroids in 3D gels within microfluidic devices containing human microvascular networks (MVNs, Figure 1a), which self-assemble over 7 days from human endothelial cells (ECs) and the same FBs used to form the tumoroids. Gene expression analysis of therapeutic target receptors on TCs in tumoroid MVN devices confirmed high ER and PR in MCF7, high HER2 in SKBR3, and high EGFR in MDA-MB-468 in the MVN devices (Figure 1b). EGFR gene expression significantly increases for all TCs isolated from the MVN devices compared to the same TCs cultured in 2D well plates, possibly due to signaling from additional cell types in the MVN devices. HER2 expression also increases approximately 10-fold for the MCF7 cells. As a result, measurable levels of EGFR and HER2 proteins are found in the microenvironment of all three tumoroids (Figure 1c). Expression of those receptors is principally co-localized with the TCs (Figure 1d), yet sectioning of tumoroids in MVN devices revealed a cytokeratin-rich core of dead TCs and, possibly, FBs where receptor expression is lost (Figure 1d, Supplementary Figure 1a). The tumoroid dead cores may be the result of hypoxia-induced necrosis, as previously observed in tumors *in vivo* and TC aggregates larger than 500 µm^25^, and as suggested by high levels of HIF-1α expression in the tumoroids (Supplementary Figure 1b). These observations cumulatively show that enhanced pathophysiological receptor expression and localization can be recapitulated in the vascularized tumoroid models compared to more simplistic 2D TC mono-culture models.

**Figure 1.**
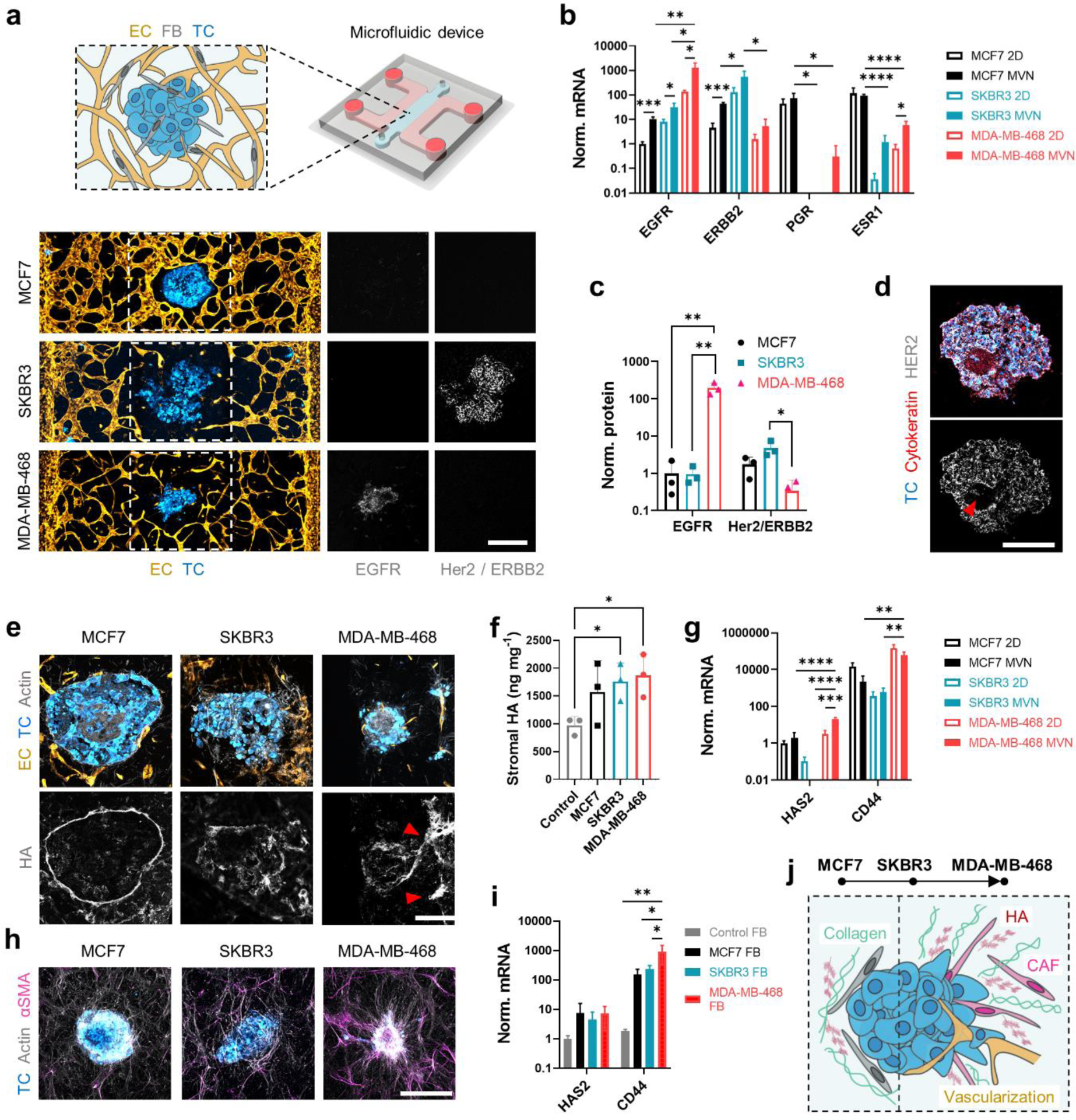
Tumoroids assembled from breast cancer lines differently remodel their surrounding stroma. (a) Schematic diagram (top) and projected confocal microscopy images (bottom) of tumoroids in the MVN devices. Expression of target receptors is confirmed by immunofluorescence. The scale bar is 500 µm. (b) Gene expression of target receptors for TCs cultured in 2D or collected from MVN devices; n = 3. (c) Protein expression of target receptors in 1 mm biopsies centered at the tumoroids in the MVN devices. (d) Projected confocal image of a SKBR3 tumoroid cryosection showing a core (arrow) where target expression is lost. The scale bar is 500 µm. (e) Projected confocal images of tumoroid cryosections showing HA localization. The red arrows indicate HA traces left by MDA-MB-468 cells migrating out of the tumoroids. The scale bar is 200 µm. (f) Quantification of stromal HA concentration in 1 mm biopsies centered at the tumoroids in the MVN devices. (g) Gene expression of HA-associated proteins for TCs cultured in 2D or collected from MVN devices; n = 3. (h) Projected confocal images of αSMA in cancer-associated FBs in the tumoroid microenvironments. The scale bar is 500 µm. (i) Gene expression of HA-associated proteins for FBs collected from MVN devices; n = 3. (j) Tumoroid microenvironments and their representation of the progression of breast cancer desmoplasia: from MCF7, where HA and collagen form a dense layer around the tumoroids, to MDA-MB-468, where HA and collagen expression increases together with the presence of cancer-associated FBs and tumoroid vascularization. Significance assessed by one-way ANOVA after confirming normal distribution of the data; in (b), (g), significance is shown only between 2D and MVNs for each TC, and between TCs in MVN; p < 0.05 *, p < 0.01 **, p < 0.001 ***, p < 0.0001 ****.

The additional complexity of the vascularized tumoroid models extends to the ways in which TCs interact with and remodel their microenvironment. MCF7 cells remain tightly packed in the tumoroids, consistent with their high expression of the epithelial junction marker E-cadherin (Supplementary Figure 1c), while the more mesenchymal SKBR3 and MDA-MB-468 cells migrate out of the tumoroids to invade the stroma (Figure 1a, e). Increased hyaluronic acid (HA) concentration is a hallmark of breast cancer desmoplasia and is associated with poor prognosis^26^. HA deposition around the tumoroids is higher compared to the surrounding microenvironment (Figure 1e). Yet, HA localization varies between tumoroids: HA forms a dense layer around MCF7 tumoroids, whereas it becomes progressively more diffuse and eventually localized with migrating TCs in SKBR3 and MDA-MB-468 tumoroids (Figure 1e). We sought to understand whether the increased HA concentration measured in the tumoroid microenvironments (Figure 1f) is a result of TC production. Gene expression analysis revealed higher production of high molecular weight HA (encoded by the HAS2 gene^27^) and expression of the HA receptor CD44 by MDA-MB-468 cells compared to the other TCs (Figure 1g). CD44 is an important adhesion molecule that TCs use to invade the stroma^26^, and its higher expression in MDA-MB-468 cells is consistent with their migratory phenotype. Interestingly, HAS2 expression in MDA-MB-468 cells significantly increases in the MVN devices compared to 2D mono-culture, again pointing to the importance of complex microenvironmental cues in modelling solid breast tumors. At the same time, SKBR3 cells have the lowest expression of HAS2 and CD44 among the TCs, and show increased expression for HA-degrading hyaluronidase 1 (HYAL1, Supplementary Figure 2a), suggesting lower HA deposition by those TCs, and the presence of an additional source of HA in their microenvironment.

Through an altered fibrotic phenotype that develops as a result of signaling in the tumor microenvironment, cancer-associated FBs (CAFs) produce aberrant amounts of HA and collagen I that promote increased TC migration and tumor proliferation^5^. Analysis by immunofluorescence of a key marker for CAFs^28^, alpha smooth muscle actin (αSMA), revealed an increasing presence of CAFs in the MCF7, SKBR3, and MDA-MB-468 tumoroid microenvironments, in respective order (Figure 1h). The larger number of CAFs in the SKBR3 and MDA-MB-468 tumoroids may be the result of increased expression of fibroblast growth factor (FGF) by those TCs^29^ (Supplementary Figure 1c). We sought to understand the role of FBs in HA production within the MVN devices, and found increased expression of CD44 (Figure 1i) and hyaluronidase 2 (HYAL2, Supplementary Figure 2b), a marker of HA metabolism^30^, in FBs in the vicinity (1 mm) of the tumoroids compared to control FBs in MVN devices without tumoroids. FBs appear more densely associated with the MDA-MB-468 tumoroids (Supplementary Figure 2c), for which we also observed increased collagen I deposition. Remarkably, collagen I becomes aligned perpendicular to the tumoroid surface in the MDA-MB-468 tumoroids, as seen through the progression of breast cancer desmoplasia to form migration tracks for TCs to invade the stroma^4^.

Overall, these results show that different tumoroids can alter their surrounding microenvironment in drastically different ways. In addition, despite the relative simplicity of TC lines compared to primary TCs, the tumoroids tested are capable of building complex microenvironments in the MVN devices that mimic key aspects of the progression of breast cancer desmoplasia^4^. Specifically, the MCF7 tumoroids appear to be representative of an early disease stage, where TCs still possess a primarily epithelial phenotype and a fibrotic ECM ‘sheath’ forms around the tumoroids that is associated with restricted TC migration and invasion; the MDA-MB-468 tumoroids, instead, may be representative of a later disease stage, whereby TCs aggressively invade the tumoroid microenvironment through a denser ECM deposited by both TCs and CAFs, with SKBR3 tumoroids somewhat in between these two extremes (Figure 1j). MDA-MB-468 tumoroids also recapitulate an additional feature of disease progression in that they often become vascularized by the surrounding MVNs (Supplementary Figure 3). We next assessed whether these vascular and ECM changes result in differences between drug distribution to the tumoroids.

### 2.2 Loss of vascular hyaluronic acid contributes to increased tumoroid interstitial fluid pressure

We have previously shown that MVNs in the vicinity of TC aggregates can partially lose vascular barrier function, resulting in higher permeability across the endothelium^23^. We observed the same phenomenon for the breast tumoroid models here, whereby focal leaks can be seen in the MVNs surrounding all three TC line tumoroids (Figure 2a). These localized leaks, in combination with a general loss of junctional integrity, result in increased MVN permeability in the vicinity of the SKBR3 and MDA-MB-468 tumoroids for dextran, a model large molecule (70 kDa) often used to benchmark vascular permeability^31^, and for Trastuzumab (HER2-targeting^32^) and Cetuximab (EGFR-targeting^33^), therapeutic monoclonal antibodies (mABs) used in the treatment of breast cancer. Remarkably, the near 3-fold increase in dextran permeability (2.8 × 10^-8^ cm s^-1^ in control MVNs to 8.1 × 10^-8^ cm s^-1^ in SKBR3 MVNs) is comparable with measurements in tumors *in vivo* (factor of ∼6)^31^.

**Figure 2.**
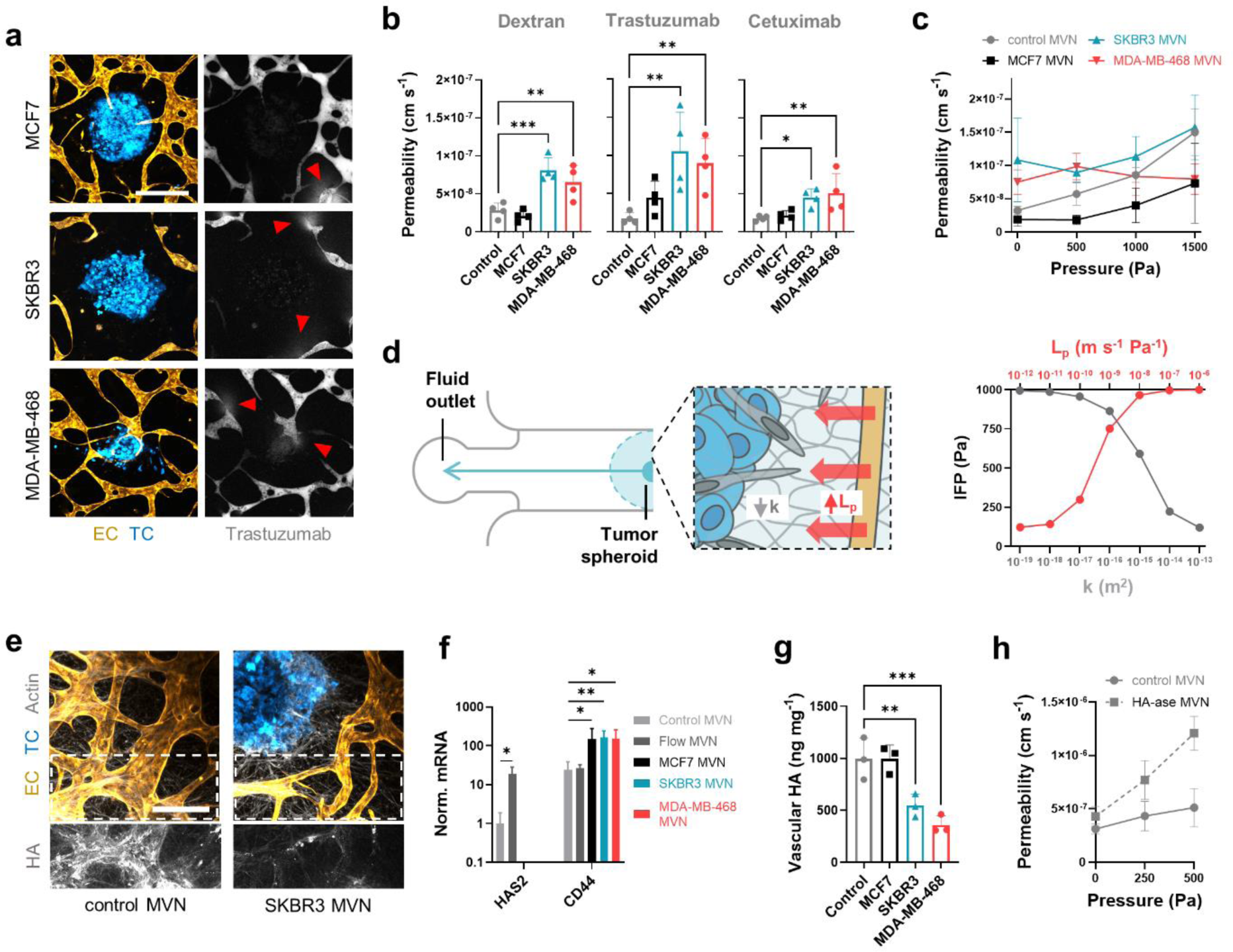
Increased vascular permeability and interstitial fluid pressure are produced by degradation of the vascular glycocalyx. (a) Confocal microscopy images of perfused tumoroid MVNs. The arrows indicate focal leaks. The scale bar is 400 µm. (b) Permeability to fluorescent dextran, Trastuzumab, and Cetuximab of tumoroid MVNs compared to control MVNs and (c) effective permeability of those MVNs as a function of applied intravascular pressure; n = 3. (d) Schematic diagram of computation model of the MVNs (left), and model results of interstitial fluid pressure, *IFP*, as a function of changes in vascular hydraulic conductivity, *L*p, and matrix permeability, *k*. (e) Confocal microscopy image of vascular HA in control and SKBR3 MVNs. (f) Gene expression of HA-associated proteins for ECs; n = 3. (g) Quantification of vascular HA concentration in 1 mm biopsies centered 2 mm from the tumoroids in the MVN devices. (h) Effective permeability of MVNs treated with HA-ase, resulting in increased filtration and hydraulic conductivity; n = 3. Significance assessed by one-way ANOVA after confirming a normal distribution of the data; p < 0.05 *, p < 0.01 **, p < 0.001 ***.

The loss of vascular barrier function appears to be incompatible with decreased drug distribution to the TME, whereby higher therapeutic molecule concentrations would be expected to reach the TCs across the leaky endothelium. However, our model allows the assessment of additional factors contributing to the impaired trans-vascular drug distribution in the tumor microenvironment to better understand this phenomenon. First, the morphology of the MVNs is altered in the vicinity of the tumoroids, as evidenced by a loss of vessel density and vessel specific surface area available for drug transport in the SKBR3 and MDA-MB-468 MVNs (Supplementary Figure 4a). Second, these morphological alterations also result in a loss of MVN perfusion capacity, whereby we observed lower tracer molecule concentrations in the vessels close to the tumoroids compared to vessels at a distance > 5 mm in the same MVN devices (by a factor of approximately 0.3 - 0.5, as measured by proxy of fluorescence intensity, Supplementary Figure 4b).

A third and key determinant of impaired drug distribution is the presence of elevated interstitial fluid pressure (IFP) in the tumoroid microenvironments. This increase in IFP near the tumoroids is observed by comparing dextran permeabilities between control and tumoroid MVNs: When control MVNs are subjected to increasing intravascular pressure, the effective dextran permeability increases with applied pressure^34^; the same trend is not observed in the tumoroid MVNs, and the resulting effective permeability at physiological intravascular pressure is comparable or lower than in control MVNs (Figure 2c). This observation indirectly demonstrates that the pressure difference across the endothelium, which drives trans-vascular fluid flow and additional molecular transport^35^, is lower in the tumoroid MVNs compared to control MVNs. This is also evidenced by the lack of interstitial fluid flow in the close vicinity of the tumoroids (Supplementary Figure 4c) and the lack of vessel diameter expansion near the tumoroids under applied intravascular pressure (Supplementary Figure 4d). Together with the lower vascular drug concentration and lower vessel surface area, the elevated IFP in the tumoroid microenvironment decreases overall drug transport across tumoroid MVNs despite the increased permeability of the vessels.

The elevated IFP in breast cancer desmoplasia hinders drug penetration and negatively correlates with cancer survival^9,31,36^. The vascularized tumoroid models present an opportunity to better understand this phenomenon and help identify therapeutic strategies that can improve drug delivery. It is well understood that increased ECM density in the desmoplastic stroma contributes to increased IFP by resisting the passage of interstitial flow via low matrix permeability, *k*^9,31^. HA is a key determinant of this contribution through its capability to bind large quantities of fluid^37^. Another, less understood contributor to elevated IFP is the leakiness of the vasculature, represented by an increased vascular hydraulic conductivity, *L*_p_. We created a 1-dimensional computational model of the tumoroid MVN devices to assess the relevant contribution of *k* and *L*_p_ to IFP in the vicinity of the tumoroids (Figure 2d, Supplementary Figure 5). Typical tumor values^38,39^ of *k* and of *L*_p_ are ≥ 10^-17^ m^2^ and ≥ 10^-11^ m s^-1^ Pa^-1^, respectively. The model results show that, starting from known values for the two factors in control MVNs (*k* ≈ 10^-12^ m^2^, *L*_p_ ≈ 10^-12^ m s^-1^ Pa^-1^)^34^, *L*_p_ changes can have a greater impact on IFP than changes in *k*, warranting additional attention to the tumoroid MVNs and the causes of their higher permeability.

Vascular *L*_p_ depends on EC junction integrity and the presence of a functional glycocalyx^35^. We did not observe changes in expression for genes associated with EC junctions between control and tumoroid MVNs (Supplementary Figure 6a). However, immunostaining of the marker ZO-1 revealed wider EC junctions near the tumoroids (Supplementary Figure 6b), consistent with previous observations of EC junctions in tumors *in vivo*^40^. HA associated with the vascular glycocalyx resists trans-vascular fluid flow as well as the passage of macromolecules^35,41^. We made a key observation that the vascular glycocalyx in the vicinity of the tumoroids is severely degraded in terms of HA concentration (Figure 2e). Strikingly, gene expression analysis for HAS2 in ECs, which encodes glycocalyx-associated HA^42^, shows repression of the gene near the tumoroids (Figure 2f). At the same time, HYAL2 expression increases (Supplementary Figure 6c) and so does CD44 expression, as previously seen for ECs in the TME^43^. Vascular glycocalyx degradation was further confirmed by a severe decrease in HA concentrations in the periphery of SKBR3 and MDA-MB-468 tumoroids (between 0.5 mm and 1.5 mm from the tumoroids, Figure 2g). Degradation of vascular HA in tumoroid MVNs also results in unchanged dextran permeability in SKBR3 MVNs after treatment of the MVNs with hyaluronidase (HA-ase), while the same treatment increases permeability in control MVNs (Supplementary Figure 6d). Crucially, HA-ase treatment of control MVNs increases *L*_p_ by one order of magnitude, from approximately 10^-12^ m s^-1^ Pa^-1^ to 10^-11^ m s^-1^ Pa^-1^ (Figure 2h).

These results confirm that vascular HA plays an important role in maintaining a low *L*_p_ in control MVNs, and that loss of vascular HA in tumoroid MVNs is associated with a concurrent increase in *L*_p_, therefore in IFP. Despite its only partial role in elevated IFP, loss of vascular HA may be used as a marker for cancer-associated vascular dysfunction. For this reason, we next assessed possible causes for this change in the tumoroid MVNs.

### 2.3 Inhibition of IL8 restores endothelial barrier function and enhances drug penetration

Vascular dysfunction in the tumoroid MVNs likely results from mechanical and/or biochemical cues specific to the aberrant microenvironments. Contact with a denser, thus stiffer, matrix can disrupt EC junctions^44^. Additionally, the lack of mechanical cues from flow in poorly perfusable tumoroid MVNs may decrease glycocalyx expression^45^. While we observed increased HAS2 expression for MVNs subjected to physiological vascular flow for 48 hours (Figure 2f) and increased vascular HA concentration (Supplementary Figure 6e), consistent with a decreased MVN permeability to dextran (4.1 × 10^-9^ cm s^-1^ compared to 1.1 × 10^-8^ cm s^-1^ in static conditions, Supplementary Figure 6d), we have previously observed functional vascular glycocalyx expression even under static MVN culture conditions^46,47^, as confirmed here by measurable expression levels for HAS2. In addition, no change in permeability or vascular HA was observed in the vicinity of the tumoroids irrespective of flow (Supplementary Figure 6d, e). Thus, impaired vascular flow in the TME may only be partially responsible for the degradation of the vascular glycocalyx. Otherwise, pro-inflammatory cytokines like tumor necrosis factor alpha (TNFα) have been previously shown to induce rapid glycocalyx shedding in ECs^48^. We, therefore, characterized the pro-inflammatory milieu found in the tumoroid MVNs to explore its role in the loss of vascular HA.

Cytokine levels in TC condition media were first assessed with a broad pro-inflammatory cytokine array (Supplementary Figure 7a). MCF7 tumoroids appear to be less disruptive to MVN function compared to SKBR3 and MDA-MB-468 tumoroids. We identified five cytokines for which concentration is two-fold higher or more for SKBR3 and MDA-MB-468 relative to MCF7 TCs: interleukin 8 and 12 (IL8, IL12), TNFα, and chemokine ligands 2 and 4 (CCL2, CCL4). Subsequent quantification of these cytokines in the lysed tumoroid microenvironments revealed higher concentrations of IL8 in both SKBR3 and MDA-MB-468 tumoroids (14 ng mL^-1^ and 35 ng mL^-1^ higher than control MVNs, respectively, Figure 3a). MDA-MB-468 tumoroids also present higher concentrations of IL12, TNFα, and CCL4, consistent with the more developed pro-inflammatory milieu expected in later stages of breast cancer desmoplasia^49^. These increased cytokine concentrations were local to the tumoroids, as analysis of supernatant collected from the MVN devices revealed much smaller changes from control MVNs (Supplementary Figure 7b).

**Figure 3.**
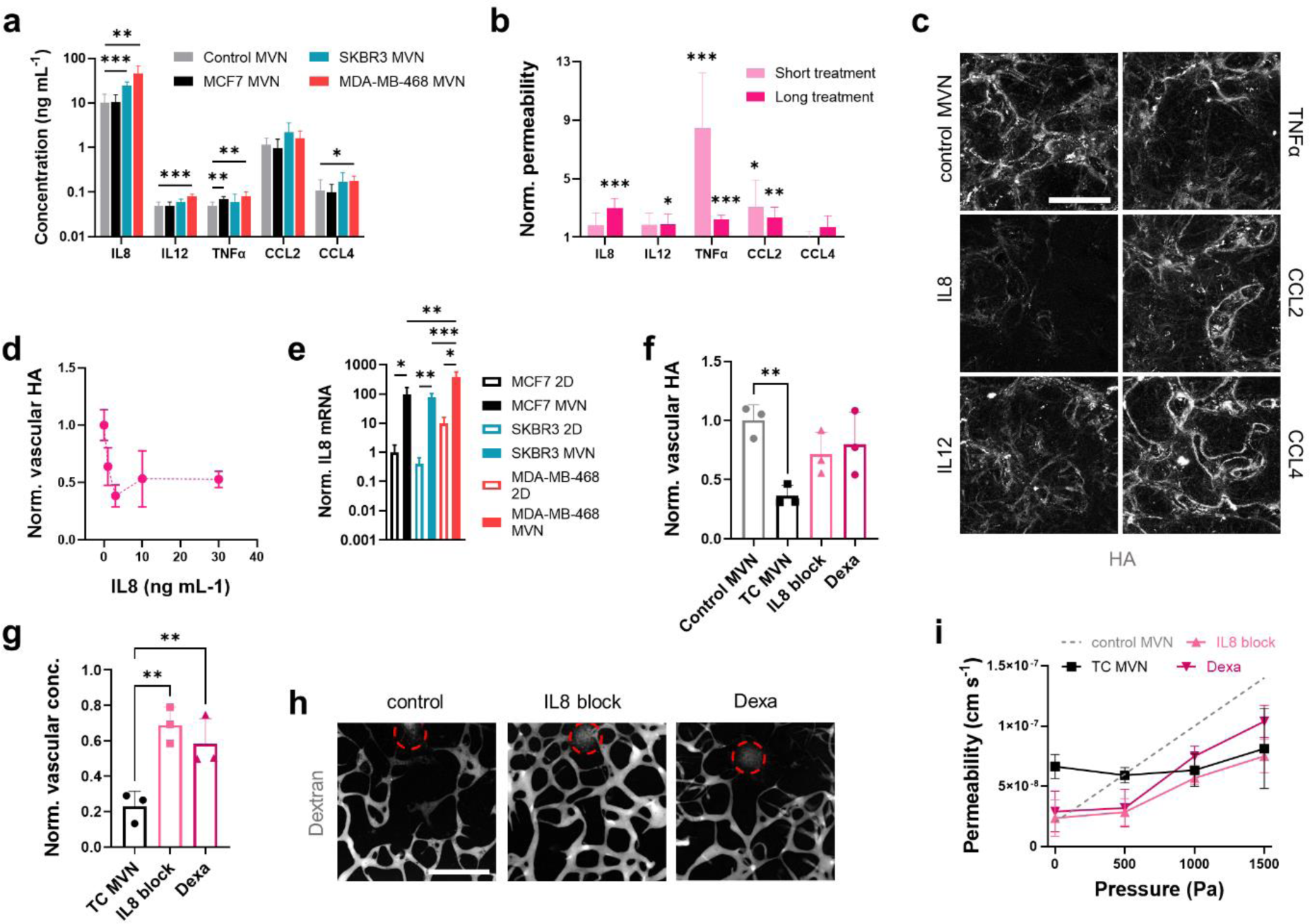
Pro-inflammatory cytokines in the tumoroid microenvironments contribute to loss of vascular glycocalyx. (a) Quantification of pro-inflammatory cytokines in 1 mm biopsies centered at tumoroids in the MVN devices; n = 3. (b) MVN permeability after short (15 min) and long (12 hours) exposure to pro-inflammatory cytokines, normalized to untreated MVN permeability; n = 3. (c) Confocal images of HA in MVNs after long exposure to the cytokines and (d) quantification of vascular HA concentration after long exposure to different concentrations of IL8; n = 3. (e) Gene expression of IL8 for TCs cultured in 2D or MVN devices; n = 3. (f) Quantification of HA concentration in 1 mm biopsies centered at MDA-MB-468 tumoroids as a result of treatment with an IL8 blocking antibody (“IL8 block”), and Dexamethasone (“Dexa”). (g) Normalized MVN concentration of dextran near the tumoroids to that at a distance > 5mm, as measured by proxy of fluorescence intensity, and (h) representative images of perfused MVNs in the vicinity of treated MDA-MB-468 tumoroids (tumoroid cores in circles). The scale bar is 500 µm. (i) Effective permeability of MDA-MB-468 MVNs with intravascular pressure as a result of the different treatments; n = 3. Significance assessed by one-way ANOVA after confirming a normal distribution of the data; in (e), significance is plotted only between 2D and MVNs for each TC; p < 0.05 *, p < 0.01 **, p < 0.001 ***.

We next set out to establish whether these cytokines directly contribute to vascular HA degradation. Recombinant versions of the five cytokines were perfused through control MVNs at a concentration of 5 ng mL^-1^ to assess changes in permeability after short (< 15 min) and long (12 hours) exposure times. TNFα and CCL2 increased MVN permeability to dextran at short times, while significant long-term changes were produced by IL8, IL12, TNFα, and CCL2, with IL8 showing the largest increase in permeability (1.9 × 10^-8^ cm s^-1^ to 5.7 × 10^-8^ cm s^-1^, Figure 3b). End-point analysis of vascular HA concentration by immunofluorescence showed significant degradation by CCL2 at short times and IL8 at long times (Figure 3c, Supplementary Figure 7c). Exposure of control MVNs to IL8 confirmed loss of vascular HA at concentrations as small as 1 ng mL^-1^ (Figure 3d). Interestingly, IL8 gene expression increases for all TCs in the MVN devices (Figure 3e, Supplementary Figure 7d). IL8 has angiogenic effects^50^, and its expression in TCs may be enhanced by paracrine signaling from ECs in the TME. Overall, these results point to IL8 as a candidate target cytokine to prevent vascular HA degradation, and subsequent loss of vascular barrier function.

We tested this hypothesis by treating MDA-MB-468 tumoroid MVNs with molecules directed against IL8: an IL8-blocking mAB, and the broad anti-inflammatory small molecule Dexamethasone. All treatments were administered in the MVN devices over 4 days. We first assessed changes in vascular HA concentrations, finding that both the IL8-blocking mAB and Dexamethasone aid in recovery of vascular HA expression to levels not significantly different from control MVNs (0.7 and 0.8 of controls, respectively, Figure 3f). The two molecules also improve MVN perfusion near the tumoroids (intravascular concentration measured as 0.7 and 0.6 of distant controls for IL8-blocking mAB and Dexamethasone, respectively, Figure 3g, h). The treatments decreased tumoroid MVN dextran permeability to values comparable to control MVNs (2.3 × 10^-8^ cm s^-1^ and 2.9 × 10^-8^ cm s^-1^ for IL8-blocking mAB and Dexamethasone, respectively, Figure 3i) and increased effective permeability with applied intravascular pressure, indicating a decrease in vascular *L*_p_, hence a decrease in IFP. Nevertheless, the lower effective permeabilities measured in the treated tumoroid MVNs compared to healthy controls confirm the continued presence of a low matrix *k* in the tumoroid microenvironments.

These results identify IL8 as an effective target to aid recovery of the vascular glycocalyx as a way of increasing drug penetration and distribution in breast tumors. Importantly, the results also suggest that targeting pathophysiological mechanisms affecting the vascular ECM can provide a different strategy to normalize the desmoplastic TME other than targeting the stromal ECM. To compare the effects of these two therapeutic strategies, we next assessed potential treatments to degrade stromal HA.

### 2.4 Desmoplastic stroma normalization strategies can be ranked for different tumoroid models

We treated, again, MDA-MB-468 tumoroids in MVNs for four days, using HA-ase, a CD44-blocking mAB, and a TGFβ-blocking mAB. While HA-ase and the CD44-blocking mAB target stromal HA directly by degrading it or blocking TCs and FBs from binding to it, respectively, the TGFβ-blocking mAB indirectly targets HA by depriving FBs in the tumoroids of TGFβ, a key stimulant of stromal HA production^51^. Similar to the treatments directed against IL8, blocking of CD44 and TGFβ in the tumoroid stroma decreased MVN permeability (2.9 × 10^-8^ cm s^-1^ for CD44, 3.8 × 10^-8^ cm s^-1^ for TGFβ, Figure 4a) and IFP, as seen by an increase in effective MVN permeability with applied intravascular pressure. This effect was particularly pronounced for the CD44-blocking treatment, whereby effective permeability values were only slightly lower than for control healthy MVNs. HA-ase, instead, did not lower permeability (5.9 × 10^-8^ cm s^-1^), but rather increased effective permeability under intravascular pressure to levels higher than control (Supplementary Figure 7e), likely the result of further degradation of vascular HA together with stromal HA. On average, all three treatments improved MVN perfusion near the tumoroids, with CD44-blocking showing the only significant increase in vascular molecular tracer concentration (to approximately 0.6 of the distant concentration levels, Figure 4b, c). These results indicate that targeting the tumoroid stromal ECM can improve vascular function, as previously explained by a decrease in vessel constriction by the dense ECM^52,53^. In particular, the decrease in tumoroid IFP, hence the increase in vascular filtration, appears more significant than what was observed when targeting the vascular ECM, as likely explained by the additional increase in matrix permeability *k* when stromal HA is degraded.

**Figure 4.**
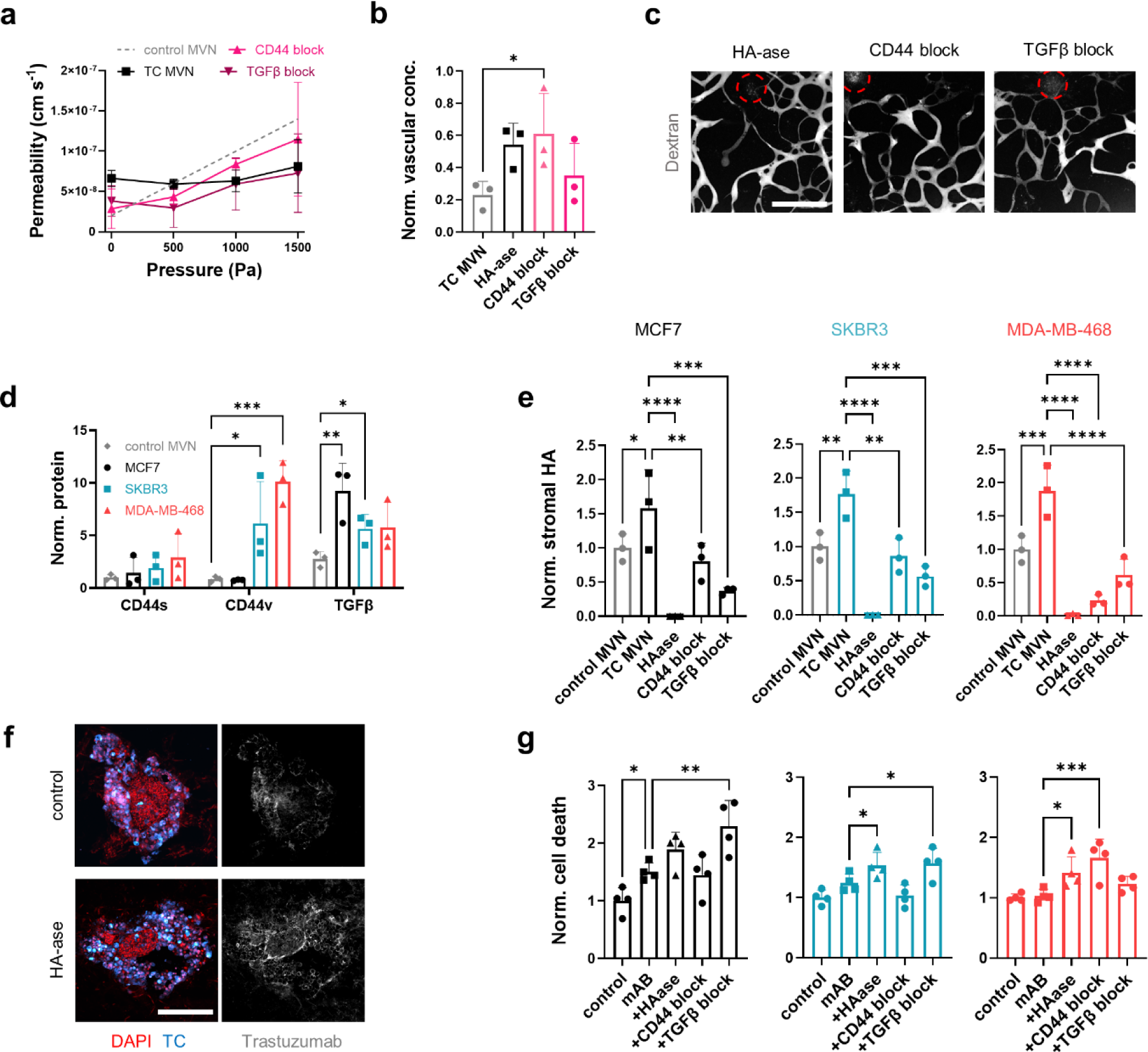
Degradation of stromal HA increases drug penetration and tumor cell death in the tumoroids. (a) Effective permeability of MDA-MB-468 MVNs with intravascular pressure as a result of different treatments targeting stromal HA; n = 3. (b) Normalized MVN concentration of dextran near the tumoroids to that at a distance > 5mm, as measured by proxy of fluorescence intensity, as a result of treatment with HA-ase, a CD44 blocking antibody (“CD44 block”), and a TGFβ blocking antibody (“TGFβ block”), and (c) representative confocal images of treated tumoroid (circles) MVNs perfused with dextran. (d) Quantification of CD44 isoforms and TGFβ in 1 mm biopsies centered at tumoroids in the MVN devices, and (e) quantification of stromal HA concentrations in the tumoroids as a result of the stromal HA-targeting treatments. (f) Confocal microscopy images of SKBR3 tumoroid cryosections showing increased Trastuzumab penetration after HA degradation. (g) Cell death in the tumoroids as a function of combined treatment with Trastuzumab + Cetuximab (“mAB”) with different targeting strategies for stromal HA, as measured by proxy of fluorescent SYTOX intensity. Significance assessed by one-way ANOVA after confirming a normal distribution of the data; p < 0.05 *, p < 0.01 **, p < 0.001 ***, p < 0.0001 ****.

Because the three TC line tumoroids show different expression and localization of HA, we next assessed how the treatments can affect stromal HA concentrations in the different tumoroids. We found increased protein expression of CD44v, the CD44 isoform associated with TC invasion^47,54^, in the SKBR3 and, more pronouncedly, MDA-MB-468 tumoroids (as opposed to the more benign isoform CD44s, which does not change from controls MVNs), and TGFβ in all three tumoroids, though highest in MCF7 (Figure 4d). Likely as a result of the different target concentrations, the treatments affected the tumoroids differently: All three treatments successfully and severely decreased stromal HA concentrations in the tumoroids, although the effect was greater by blocking TGFβ in the MCF7 and SKBR3 tumoroids and by blocking CD44 in the MDA-MB-468 tumoroids (Figure 4e). The greatest effect was achieved by HA-ase, which fully degraded HA in all three tumoroid types.

We further hypothesized that degradation of stromal HA can lead to increased drug distribution to TCs. To test this, the tumoroid MVNs were treated for four days with Trastuzumab and Cetuximab, a model drug combination with expected cytotoxic effects^32,33^ to compare drug effectiveness in the different tumoroids, at a concentration of 20 µg mL^-1^ to match expected levels in circulation^34^. Preliminary data showed significant TC death in tumoroids in well plates at concentrations as low as 2 µg mL^-1^ when assessed by fluorescence imaging of a cell death marker (Supplementary Figure 8a, b). We used the same method to evaluate cell death as a result of co-perfusion with the molecules targeting stromal HA in the MVN devices. Sections of the fixed tumoroids post-treatment revealed increased drug penetration compared to controls (Figure 4f), validating our hypothesis that stromal HA degradation increases interstitial drug transport. Despite the relatively high mAB concentration perfused, significant cell death was observed only in the MCF7 tumoroids (Figure 4g), confirming the negative effect of impaired vascular perfusion and increased ECM density on drug distribution to the tumoroids. Co-treatment with HA-ase of SKBR3 and MDA-MB-468 tumoroids increases TC death, and so does co-treatment with the TGFβ-blocking mAB for MCF7 and SKBR3 tumoroids, and with the CD44-blocking mAB for MDA-MB-468 tumoroids. Measurement of size change in the tumoroids was less conclusive, as a significant decrease in size over the four days can only by seen with co-treatment with HA-ase (Supplementary Figure 8c), likely due to its particularly efficient degradation of stromal HA.

Overall, these results confirm our hypothesis that stromal HA degradation in the tumoroid microenvironments correlates with increased drug efficacy due to improved penetration. The results also showcase the capability of the models to assess relative impact for different therapeutic strategies in different tumoroid models. We next applied this capability to rank therapeutic strategies in vascularized tumoroids assembled from patient-derived TCs.

### 2.5 Personalized tumoroid models pinpoint effective desmoplasia normalization strategies

We formed tumoroids with TCs from two breast cancer patients and cultured them within MVN devices (Figure 5a). Both patient-derived tumoroids become more highly vascularized by the MVNs compared to TC line tumoroids. The effect is particularly pronounced for Patient 2, for which the perfused dextran concentration near the tumoroids is not different from that in distant MVN vessels (Supplementary Figure 9a). TCs from Patient 2 are also more invasive and seen to actively colonize the tumoroid MVNs (Figure 5a), consistent with the increased aggressiveness of the particular breast cancer sub-type^24^ (basal, compared to luminal for Patient 1). The morphology of the MVNs is similar between the two patient TCs, but the patient tumoroid MVNs possess a larger volume fraction and specific surface area compared to control MVNs (Supplementary Figure 9b). Vascular barrier function of the patient tumoroid MVNs is altered to a greater extent than the SKBR3 and MDA-MB-468 tumoroids, with increased permeability (1.5 × 10^-7^ cm s^-1^ and 9.8 × 10^-8^ cm s^-1^ for Patient 1 and Patient 2, respectively) and a lack of a clear trend with applied intravascular pressure, confirming an increase in IFP even for these tumoroids (Figure 5b). These results further confirm that our models can display a wide range of breast cancer heterogeneity between different disease types and patients.

**Figure 5.**
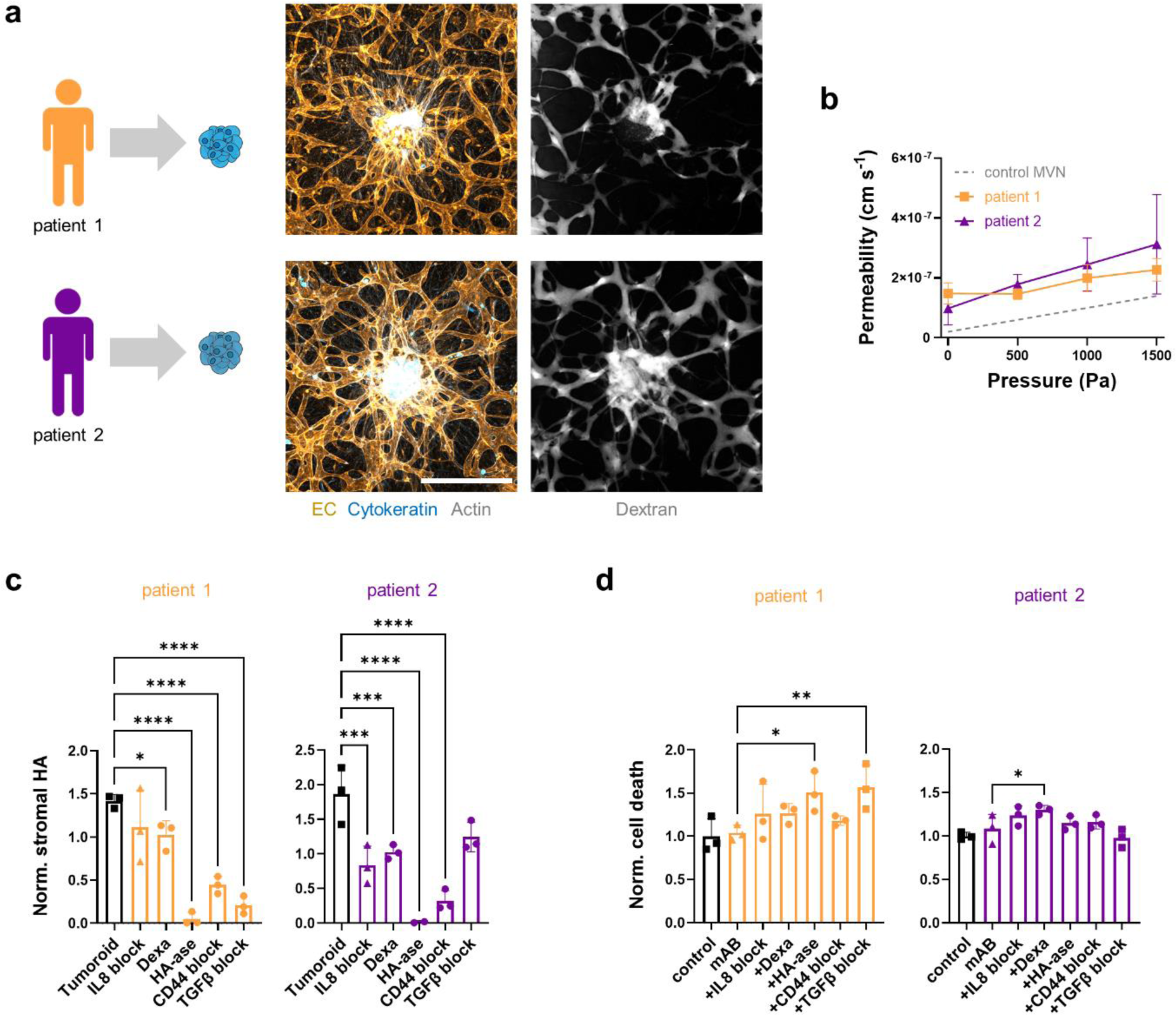
Vascularized tumoroids from primary breast cancer cells respond differently to treatments normalizing the stromal and vascular extra-cellular matrix. (a) Projected confocal microscopy images of patient-derived tumoroids show high levels of vascularization and different levels of cancer vascular colonization. (b) Effective permeability of patient-derived tumoroid MVNs with intravascular pressure; n = 3. (c) Quantification of stromal HA concentrations in the tumoroids as a result of stromal and vascular HA-targeting treatments, and (d) resulting cell death in the tumoroids when the treatments are administered in combination with Trastuzumab and Cetuximab, as measured by proxy of fluorescent SYTOX intensity. Significance assessed by one-way ANOVA after confirming a normal distribution of the data; p < 0.05 *, p < 0.01 **, p < 0.001 ***, p < 0.0001 ****.

Both patient-derived tumoroids show higher stromal HA concentrations than control MVNs (1410 ng mg^-1^ and 1854 ng mg^-1^ for Patient 1 and Patient 2, respectively, compared to 994 ng mg^-1^). We tested the treatment regimens described above to attempt normalization of the tumoroid microenvironments, including those treatments rebuilding vascular HA and those degrading stromal HA (Figure 5c). All treatments produce an overall decrease in stromal HA concentration, although not significant for the IL8-blocking mAB in Patient 1 tumoroids and the TGFβ-blocking mAB in Patient 2 tumoroids. Aside from HA-ase, which virtually obliterates HA in the tumoroids, the decrease in HA concentration is greatest when blocking TGFβ in Patient 1 tumoroids and blocking CD44 in Patient 2 tumoroids. Remarkably, while the IL8 blocking mAB and Dexamethasone decrease overall HA concentrations less than the other treatments, they produce levels comparable to MVN controls, possibly indicating a greater extent of ECM normalization by targeting both the vascular and stromal HA. Most importantly, these results show that even when assembled from primary TCs, the vascularized tumoroid models are capable of capturing variations in the effectiveness of different treatments in normalizing the desmoplastic microenvironments.

Finally, we assessed the efficacy of the different treatments in producing changes in TC death when used in combination with Trastuzumab and Cetuximab, as previously done with the TC line tumoroids. We found that Patient 1 tumoroids are more susceptible to treatment with the therapeutic mABs compared to Patient 2 tumoroids (Figure 5d), likely the result of differential expression of target receptors between the two TC types (HER2 expression confirmed by the supplier only for Patient 1 TCs). For Patient 1, cell death increases significantly with combination treatment with HA-ase and the TGFβ-blocking mAB, for which HA degradation was greatest, as similarly seen in MCF7 tumoroids. Despite the overall inefficacy of Trastuzumab and Cetuximab in targeting Patient 2 tumoroids, we observed increased cell death with combination treatment of the mABs with Dexamethasone (and, although not significant, with the IL8-blocking mAB). A different treatment approach, for example by standard small molecule chemotherapy or different mAB target, may prove more effective in producing significant TC death for Patient 2. Taken together, these results confirm that patient-derived vascularized tumoroid models can help rank therapeutic strategies by effectiveness in normalizing the tumoroid microenvironment and improving delivery and efficacy of clinically relevant therapies.

## 3. DISCUSSION

Normalization of desmoplastic cancers is currently being approached therapeutically by targeting cell contractility of TCs and CAFs or their aberrant ECM deposition^10^, both with the purpose of preventing blood vessel constriction and improving drug penetration. Of these treatments, TGFβ-targeting appears particularly attractive due to the key role uncovered for this factor in stimulating desmoplastic collagen and HA deposition^55^. Our results using vascularized tumoroid models confirm that targeting TGFβ can be successful in normalizing the tumoroid microenvironments for certain patients^55^, as seen in models of early-stage breast cancer (MCF7 and SKBR3 TC lines and primary luminal TCs). At the same time, direct targeting of HA through CD44 can achieve greater normalizing responses for other patients, as seen in our results for models of advanced, aggressive breast cancer (MDA-MB-468 TC line and primary basal TCs). This observation is consistent with the identification of HA and CD44 as markers of metastatic breast cancer ^54,56^, and that their inhibition can prevent vascular colonization by TCs^47^. Importantly, our results show that the vascularized tumoroid models can capture these variations in desmoplastic ECM deposition and resulting impaired MVN perfusion, allowing the assessment of different therapeutic strategies to improve drug penetration.

Notably, using our models we have shown that HA exists in two forms in the desmoplastic microenvironment: vascular HA and stromal HA, both of which constitute biomechanical barriers to drug penetration. Vascular HA degradation in the tumoroid microenvironment may not only impair vascular barrier function, but also favor TC vascular migration and metastatic colonization^47^. Stromal HA degradation, instead, can prove beneficial to treatment by improving interstitial transport of therapeutic molecules to TCs. HA-ase, in the relatively high concentrations used here, results in complete obliteration of both vascular and stromal HA in the tumoroid microenvironments, thus in exaggerated fluid filtration that may produce edema formation *in vivo*. It is tempting to speculate that treatments that can recover vascular HA while degrading stromal HA may be particularly beneficial in normalizing the breast cancer microenvironment and prevent disease progression. We identified IL8 as a potential target to achieve both vascular and stromal HA normalization, as demonstrated by recovery of MVN perfusion and vascular barrier function after treatment by a blocking mAB or by the broad anti-inflammatory drug Dexamethasone. IL8 can increase the metastatic potential of breast TCs^57^ and IL8 serum concentrations have been shown to increase significantly in patients with advanced breast cancer, correlating negatively with patient survival^58^. Treatments targeting IL8 and the tumor pro-inflammatory milieu may find new use in normalizing breast cancer desmoplasia, as recently demonstrated by increased stromal and vessel normalization by Dexamethasone in murine breast cancer models^59^.

The heterogeneity of breast cancer desmoplasia makes patient-specific *in vitro* models uniquely positioned to pinpoint treatment strategies that are most effective in achieving a therapeutic response^15^. The MVN models allow for culture of patient-derived tumoroids in functional microvascular beds that allow perfusion of relevant therapeutic molecules under physiological vascular flow conditions. In addition, tumoroids from different parts of an excised tumor may also be tested individually to address intratumor heterogeneity. MVNs were formed here with primary human umbilical vein ECs (HUVECs) and lung FBs, which were chosen for their vasculogenic potential and robust formation of perfusable MVNs even when exposed to pro-inflammatory factors^23,47^. Future modifications of the models may include immortalized MVN cells sources with improved reproducibility^60^, or primary mammary ECs and FBs cultured under constant vascular flow^61^ for improved physiological tissue fidelity and model longevity. Patient-derived tumoroids including native immune cell populations and ECM^13^ may also be incorporated in the models to assess the efficacy of immunotherapies. Irrespective of cell sources, the demonstrated capability of these models to quantify the efficacy of different therapeutic strategies attests to the formation of a relevant pathophysiological microenvironment in the vascularized breast tumoroid models. These models may find application as personalized translational assays to inform clinical therapeutic strategies and identify those with increased probability of treatment success.

## 4. METHODS

### 4.1 Vascularized tumoroid formation

TC lines MCF7, SKBR3, and MDA-MB-468 were obtained from ATCC and cultured in Dulbecco’s Modified Eagle’s Medium (#10566016, ThermoFisher) supplemented with 10% fetal bovine serum (#12662029, ThermoFisher). All TC lines were made to express red fluorescent protein (RFP) as previously described^47^. Primary TCs were obtained from Amsbio (#MBE-F-TM and #MLE-F-TM) and cultured in Mammary Epithelial Cell Growth Medium (#C-21010, Promocell). Human umbilical vein ECs (HUVECs, wild type or GFP-expressing) from pooled donors were obtained from Angio-Proteomie (#cAP-0001 and #cAP-0001GFP) and cultured in Vasculife Endothelial Medium (#LL-0003, Lifeline) up to passage 5. Normal human lung FBs were obtained from Lonza (#CC-2512) and cultured in Fibrolife Fibroblast Medium (#LL-0011, Lifeline) up to passage 5. Tumoroids were self-assembled by co-culture of 4k TCs and 5k FBs in non-adherent 96-well plates (PrimeSurface 96M, Sbio) over 4 days without change in culture medium. MVNs incorporating tumoroids were formed as previously described^23^ in a three-channel microfluidic device (central gel channel: 3 mm × 0.5 mm × 1cm)^34,46,62^ made from polydimethylsiloxane (PDMS, Ellsworth Adhesives) bound to no. 1 glass coverslips (#48404-097, VWR). ECs (6 million mL^-1^), FBs (2 million mL^-1^), and tumoroids were co-injected with fibrin gel solution within the central channel of the microfluidic device, and cultured over 7 days through daily Vasculife medium changes in the side channels. A monolayer of ECs was seeded on the gel surfaces in the side channels on day 4 of culture, as previously described^46^.

### 4.2 MVN permeability and fluid flows

Permeability of fluorescent molecules in the MVNs was measured by confocal microscopy and image analysis with the software ImageJ as previously described^46^, using fluorescein isothiocyanate (FITC, #F4274, Sigma Aldrich), dextran (FITC-conjugated, 70 kDa, 0.1 mg mL^-1^, #FD70, Sigma Aldrich), Trastuzumab (Alexa Fluor 488-conjugated, #FAB9589G, 0.1 mg mL^-1^, R&D Systems), and Cetuximab (Alex Fluor 647-conjugated, #FAB9577R, 0.1 mg mL^-1^, R&D Systems). The permeability analysis also yields the morphological parameters measured^46^: vascular volume fraction, *V%*, specific surface area, *SSA*, and average vessel diameter, *d*. The MVNs were conditioned through vascular flow for 48 hours using a custom pump that applies a constant pressure difference across the MVNs of approximately 50 Pa^63^. Pressurization of the MVNs through the microfluidic device side channels was done as shown previously^34^ using a FlowEZ pressure regulator (Fluigent), and the effective MVN permeability for intravascular pressures up to 1500 Pa was measured as above. The increase in effective permeability of FITC was used to calculate the hydraulic conductivity of the MVNs, *L*_p_, using^64^:

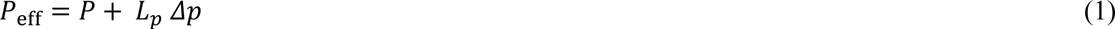

where *P* is the MVN permeability at intravascular pressure Δ*p* = 0 Pa and *P*_eff_ is the effective permeability. No endothelial reflection of the small solute and no difference in osmotic pressure across the endothelium were assumed, as demonstrated before for this system^34^. The interstitial flow resulting from MVN pressurization in the vicinity of the tumoroids was measured by fluorescence tracking of a 30 µm bleached spot, as previously described^34^.

### 4.3 Interstitial fluid pressure modeling

Computational modeling of IFP in the MVN devices was performed using the software COMSOL Multiphysics v5.4. The gel channel of the microfluidic device was described by a 1D domain divided into four subsequent zones: the tumor spheroid (A – 0 to 0.5 mm), the gel areas close to the spheroid (B – 0.5 to 3 mm) and far from it (C –3 to 5 mm), and the exit portion of the chip (D –5 to 7 mm). Only half of the chip was simulated due to symmetry, identifying the spheroid center as origin of the domain (*x* = 0, Figure 2d). Fluid flow in the gel was accounted for by the Darcy equation, and the Starling equation was used to include fluid filtration from both the MVNs and the lateral monolayers, neglecting osmotic differences^46^. The resulting equations were:

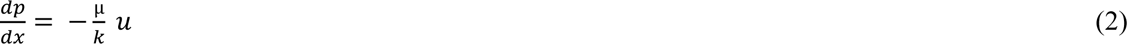

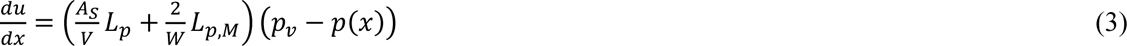

where *p* is the fluid pressure, *u* is the fluid velocity, *µ* is the fluid viscosity, *k* is the matrix permeability, *A*_S_/*V* is the ratio of the MVN lateral surface and the matrix volume, *L*_p,M_ is the hydraulic conductivity of the side monolayer, assumed one order of magnitude higher than that of the MVNs based on our previous assessments^34^, and *W* is the width of the gel channel. Boundary conditions complemented the model imposing the symmetry condition (*u*(0) = 0), and the atmospheric pressure as outlet (*p*(7mm) = 0). Parameter values are reported in Supplementary Table 1. Geometrical differences at the exit of the microfluidic device were included in the model as a change of Darcy equation parameters. The relative effect of *Lp* and *k* in the B area of the device was investigated by changing their values across different orders of magnitude and analyzing resulting fluid pressure and velocity.

### 4.4 Gene and protein expression

Gene expression was assessed by quantitative polymerase chain reaction (qPCR). Tumoroids from 4 microfluidic devices, 4 tumoroids per device, were collected by separating PDMS and glass using a scalpel to access the gel, as previously described^65^. Gel biopsy punches (1 mm diameter, #15110-10, Ted Pella) centered at each tumoroid were collected and the matrix between cells lysed with Liberase (5 mg mL^-1^, #5401135001, Sigma Aldrich) for 30 minutes. ECs (GFP), TCs (RFP), and FBs (non-labeled) were then sorted in Trizol (#15596026, Fisher Scientific) using BD FACSAria™ III Cell Sorter. Dead cells were eliminated by DAPI staining. Custom PCR plates (TaqMan 96 well-plate, ThermoFisher) were used to assess mRNA expression on a QuantStudio 12K Flex Real-Time PCR System (ThermoFisher) for the following genes: GAPDH, EGFR, ERBB2, PGR, ESR1, ABCB1, ABCG2, CXCL8, IL12A, CCL2, CCL4, TNF, VEGFA, FGF1, CDH1, HAS1, HAS2, HAS3, HYAL1, HYAL2, CD44, TJP1, CLDN5, OCLN, FCGRT, CAV1, CLTC. Gene expression was normalized to 18S endogenous control and GAPDH. Missing data values represent non-detectable signals. Protein expression was assessed in four ways: i) immunofluorescence staining of EGFR (Cetuximab, as above), HER2 (Trastuzumab, as above), cytokeratin (#M3515, Agilent), HA (HA binding protein, #385911, Sigma Aldrich), αSMA (#19245S, Cell Signaling Technology), HIF-1α (#ab8366, Abcam), Caspase-3 (#9661S, Cell Signaling Technology), COL-1 (#AF6220-SP, R&D Systems), ZO-1 (#33-9100, ThermoFisher); ii) ProteinSimple automatic western analysis, as described previously^47^, of EGFR (#MAB9577, R&D Systems), HER2 (#MAB9589, R&D Systems), CD44 (#GTX102111, GeneTex, isoform assessed at peaks CD44s: 90 kDa, and CD44v: 160 kDa), TGFβ (#MAB1835, R&D Systems); iii) Enzyme linked immunosorbent assay (ELISA) of HA (#DHYAL0, R&D Systems), normalized to total protein content, and pro-inflammatory cytokines (array, #ab134003, Abcam); iv) custom Meso Scale Discovery (MSD) assay of pro-inflammatory cytokines IL8, IL12, CCL2, CCL4, TNFα. Protein quantification was performed on single 1 mm gel biopsy punches lysed in buffer (#9803S, Cell Signaling Technologies) containing Benzonase Nuclease (#E8263, Sigma Aldrich) and Protease inhibitor cocktail (#11836170001, Sigma Aldrich). Immunostaining was performed on either intact gels fixed with 4% paraformaldehyde (#50-259-96, Fisher Scientific) within microfluidic devices, or on gels extracted from devices and embedded in Optimal cutting temperature (OCT) compound before cryosectioning on a Leica 1850 cryotome.

### 4.5 MVN and tumoroid treatments

Treatments were administered in the microfluidic devices through perfusion across the MVNs. HA-ase (150 units mL^-1^, #H3631, Sigma Aldrich) was perfused for 30 min under a transient pressure difference across the MVNs before measurement of *L*_p_. Recombinant human IL8 (#208-IL-010, R&D Systems), IL12 (#219-IL-005, R&D Systems), TNFα (#210-TA-005, R&D Systems), CCL2 (#279-MC-010, R&D Systems), CCL4 (#271-BME-010, R&D Systems) were perfused at a concentration of 5 ng mL^-1^ together with dextran for short treatment-permeability testing (< 15 min) and for 12 hrs for long treatment-testing. MVN and tumoroid treatments were administered over 4 days through daily media perfusion with an IL8 blocking antibody (20 µg mL^-1^, #MAB208, R&D Systems), Dexamethasone (5 µM, #D4902, Sigma Aldrich), HA-ase (as above), a CD44 blocking antibody (10 µg mL^-1^, 08-9407-2, American Research Products), and a TGFβ blocking antibody (20 µg mL^-1^, #MAB1835, R&D Systems). Trastuzumab (#MAB9589, R&D Systems) and Cetuximab (#MAB9577, R&D Systems) were co-perfused at a concentration of 20 µg mL^-1^, and changes in cell death in the tumoroids assessed through imaging of SYTOX Green (#S34860, ThermoFisher) on a Nikon Eclipse Ti microscope with a 4X objective. The SYTOX signal intensity was measured using ImageJ in the area co-localized with the RFP TC signal. Changes in size of the tumoroids as a result of the different treatments were measured in terms of changes in projected area, also through the RFP TC signal. For the primary, non-fluorescent TCs, the SYTOX signal intensity was measured in a circular area of diameter 1 mm from the center of the tumoroids.

### 4.6 Data interpretation and statistics

The number of technical and biological repeats is reported in each figure caption. When possible without compromising clarity, single biological repeat data points are provided in the graphs. Statistical analysis of the data was performed with the software Prism (GraphPad, version 9), and the specific statistical test used is reported in each figure caption. Mean differences with *p value* < 0.05 were taken as significant.

## ACKNOWLEDGEMENTS

We are thankful to Patricia Amarante and Diana Cha at Amgen Inc. for training and access to the ProteinSimple western and the MSD assays. We also thank the Flow Cytometry Core facility at Koch Institute for Integrative Cancer Research at MIT for technical support. G.S.O. received funding from Amgen and an American Italian Postdoctoral Fellowship. Z.W. is supported by the Ludwig Center Fund Post-Doctoral Research Fellowship and by National Cancer Institute (U01 CA214381). L.P. was supported by AIRC Investigator Grant, no. IG21479. H.T.N. was supported by a Swiss National Science Foundation Postdoctoral Fellowship (SNSF-P400PB_186779).

## AUTHORS CONTRIBUTIONS

G.S.O., C.G.K., and R.D.K. designed the study; G.S.O., K.H., Z.W., H.T.N., and M.R.G. performed the experiments; L.P. performed the computational modeling. G.S.O. wrote the first draft of the manuscript, and all authors contributed to its final form and scientific discussion.

## DATA AVAILABILITY

The datasets generated during the current study are available from the corresponding authors on reasonable request.

## SUPPLEMENTARY INFORMATION

**Supplementary Figure 1.**
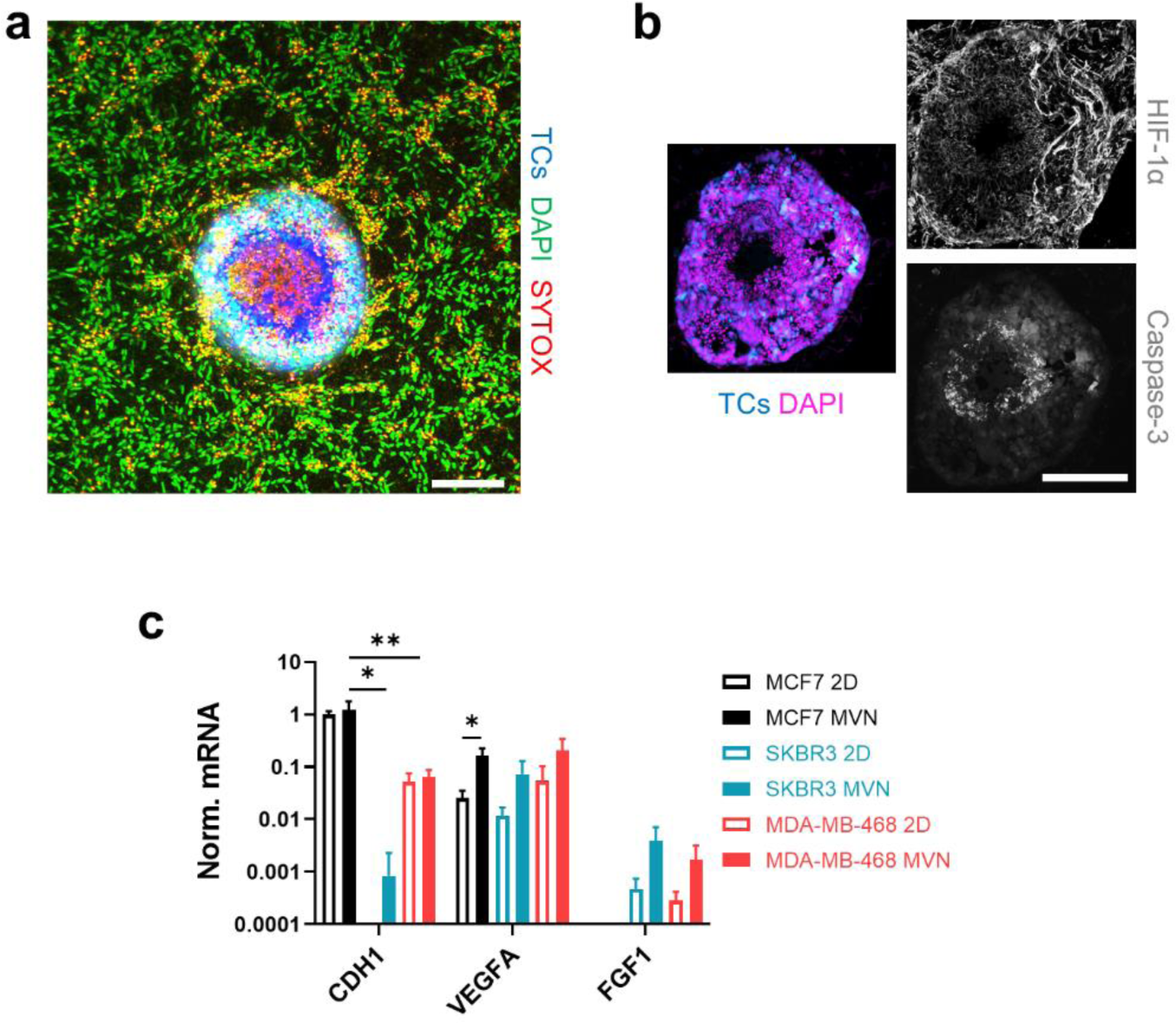
(a) Projected confocal microscopy image showing dead core (SYTOX signal) in an MCF7 tumoroid in MVN devices. The scale bar is 200 µm. (b) Confocal image of MCF7 tumoroid cryosection showing elevated HIF-1α. The scale bar is 200 µm. (c) Gene expression of TC lines cultured in 2D or MVNs; n = 3. Significance assessed by one-way ANOVA; p < 0.05 *, p < 0.01 **.

**Supplementary Figure 2.**
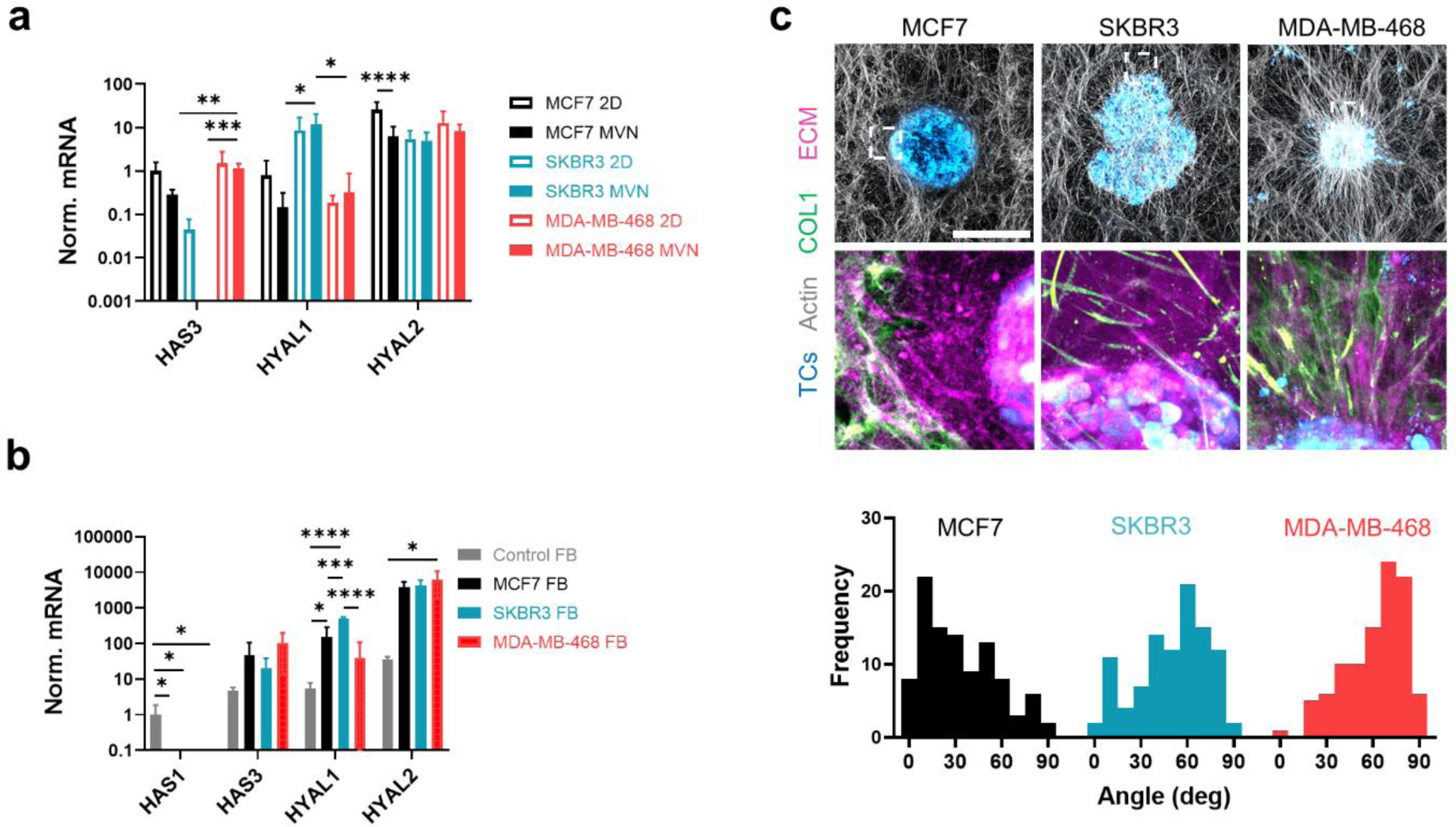
(a) Gene expression in TC lines cultured in 2D or in MVNs; n = 3. (b) Gene expression in FBs from tumoroids or control MVNs; n = 3. (c) Confocal images of collagen I in the tumoroids (top, scale bar = 500 µm), and distributions in the direction of collagen I streaks (bottom); n = 100. Significance assessed by one-way ANOVA; p < 0.05 *, p< 0.01 **, p < 0.001 ***, p < 0.0001 ****.

**Supplementary Figure 3.**
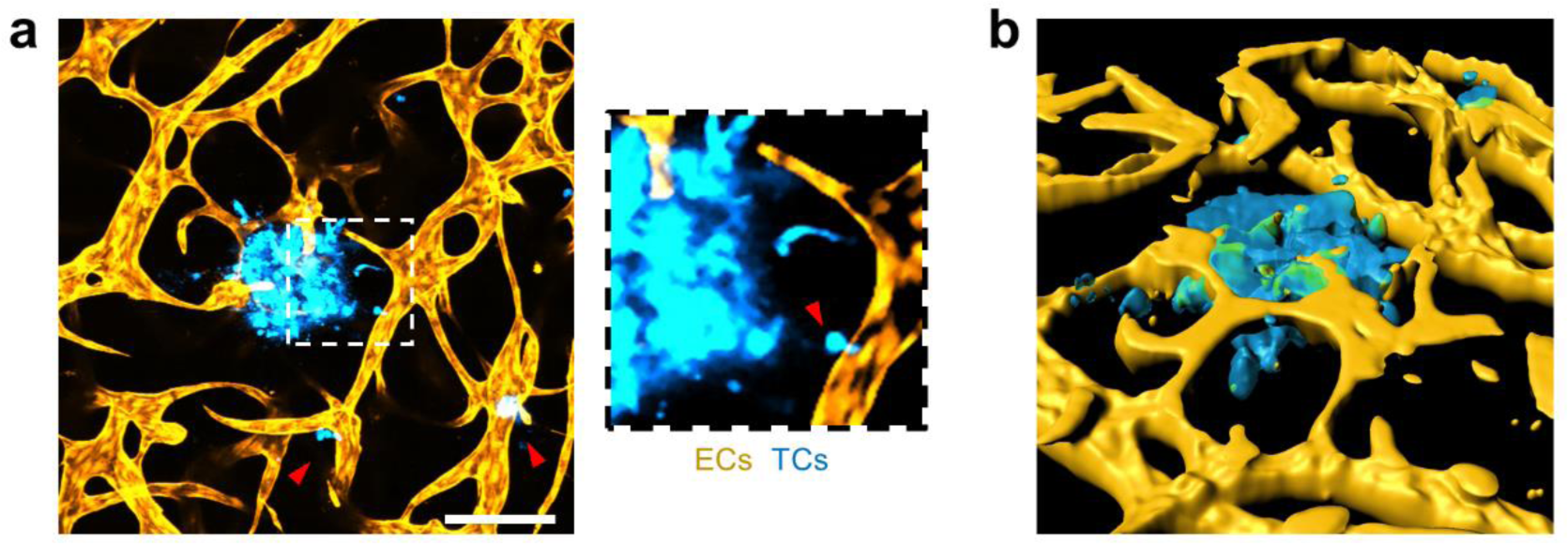
(a) Confocal image of vascularized MDA-MB-468 tumoroid (scale bar = 200 µm) and (b) its 3D reconstruction using the Imaris software showing vessel penetration into the tumoroid mass. The arrows in (a) indicate clusters of invasive TCs. The inset shows a TC intravasating into the MVNs.

**Supplementary Figure 4.**
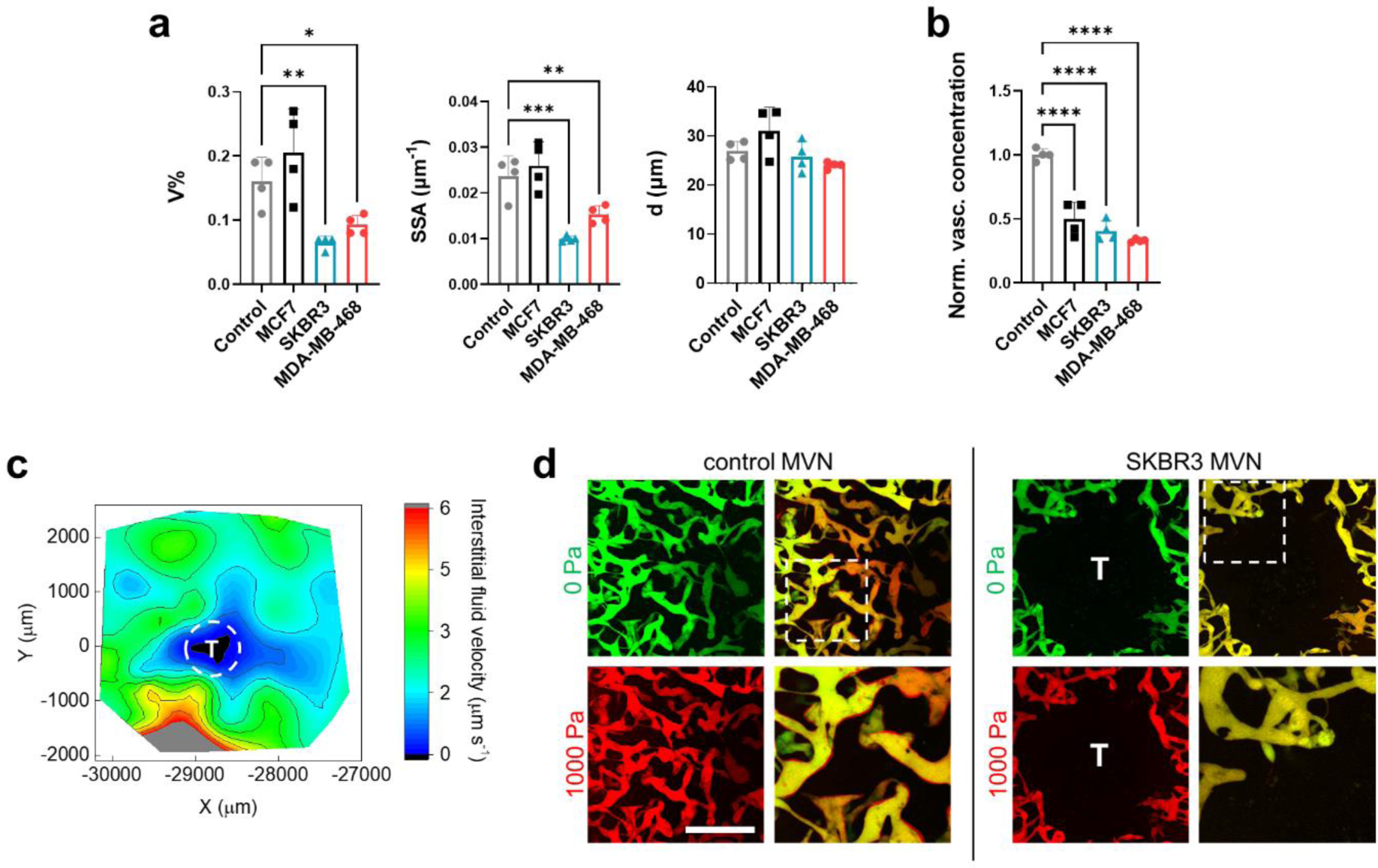
(a) Morphological characterization of control and tumoroid MVNs in terms of volume fraction, *V%*, specific surface area, *SSA*, and vessel diameter, *d*, and (b) normalized dextran concentration in those MVNs, as measured by proxy of fluorescence intensity. (c) Interstitial fluid velocity in the vicinity of a MDA-MB-468 tumoroid in the MVN device, as measured by fluorescence tracking. (d) Fluorescence microscopy images of control and SKBR3 MVNs perfused with dextran as a function of intravascular pressure (left, 0 Pa = green, 1000 Pa = red). Merged images (right) show vessel expansion due to intravascular pressure as a red outline. Expansion of the vessels can be seen in pressurized control MVNs, but not in tumoroid MVNs. The scale bar is 500 µm. Significance assessed by one-way ANOVA; p < 0.5 *, p < 0.01 **, p < 0.001 ***, p < 0.0001 ****.

**Supplementary Figure 5.**
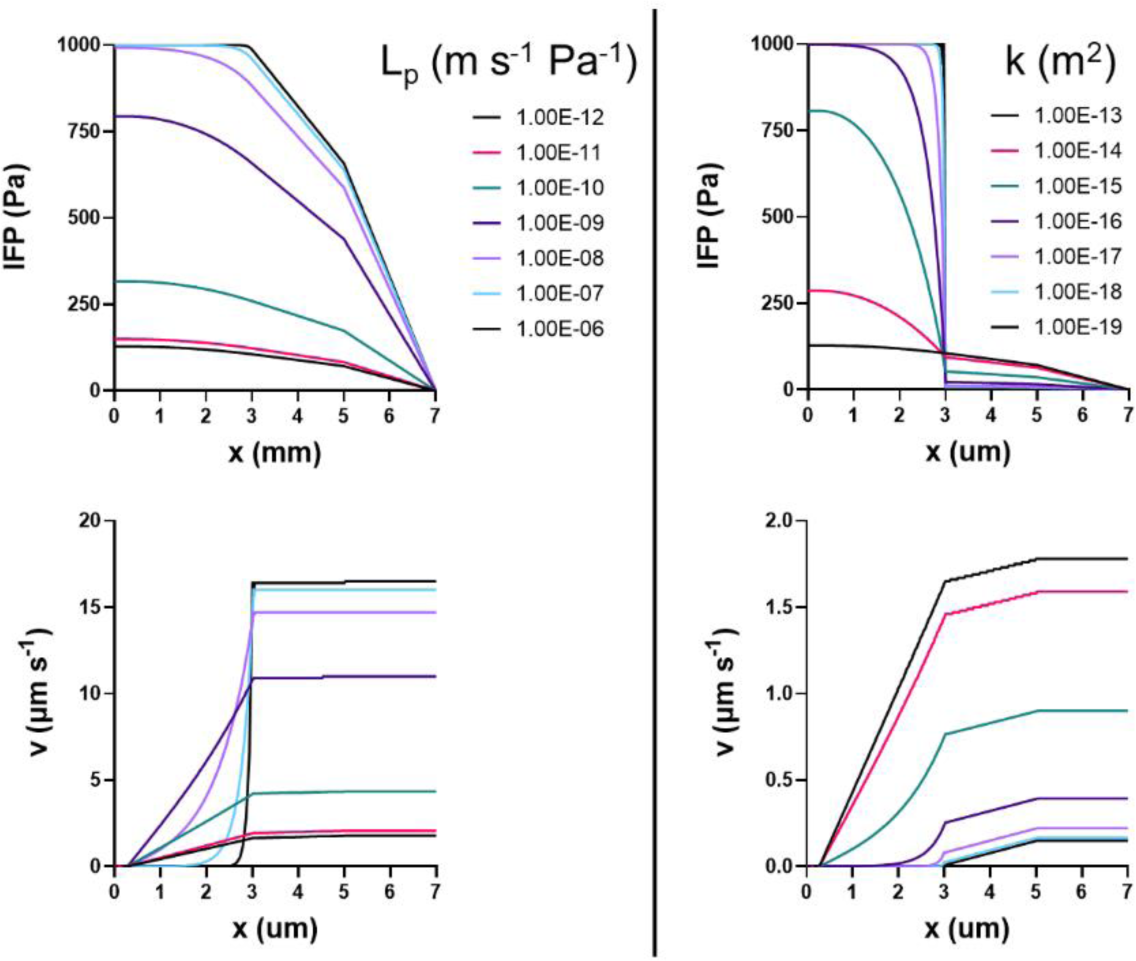
Computational model output of interstitial fluid pressure (top) and interstitial fluid velocity (bottom) as a function of distance from the tumoroids and changes in vascular hydraulic conductivity (left) or matrix permeability (right).

**Supplementary Figure 6.**
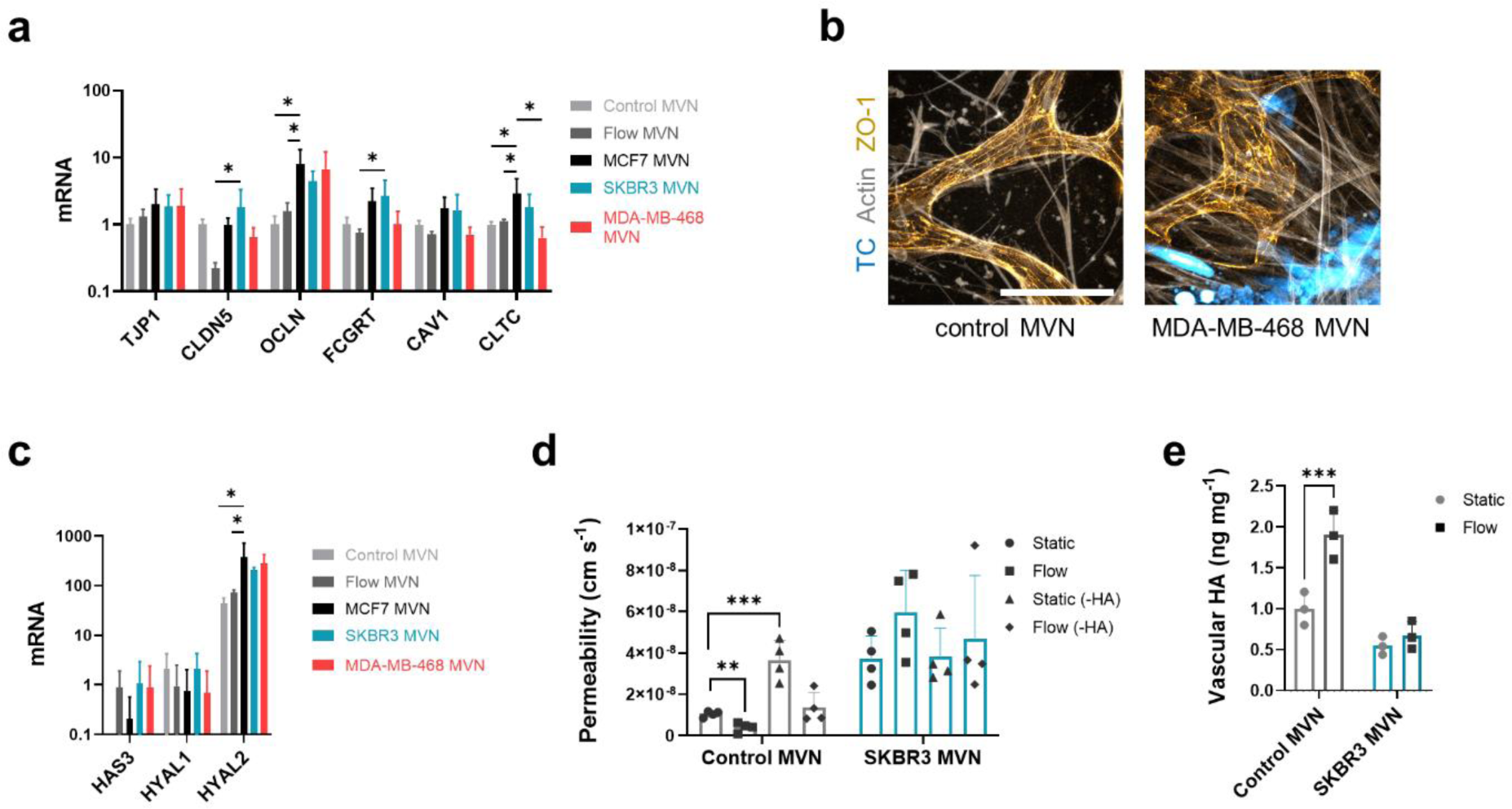
(a) Gene expression in ECs from control or tumoroid MVNs, for genes associated with vascular transport; n = 3. (b) Confocal microscopy image of tight junctions in MVNs. The scale bar is 100 µm. (c) Gene expression of ECs for genes associated with HA production; n =3. (d) HA concentration in control and SKBR3 MVNs cultured statically or under continuous vascular perfusion for 48 hrs. (e) Resulting MVN permeability as a function of flow condition, proximity to the tumoroid, and HA degradation by HA-ase. Significance assessed by one-way ANOVA; p < 0.5 *, p < 0.01 **, p < 0.001 ***, p < 0.0001 ****.

**Supplementary Figure 7.**
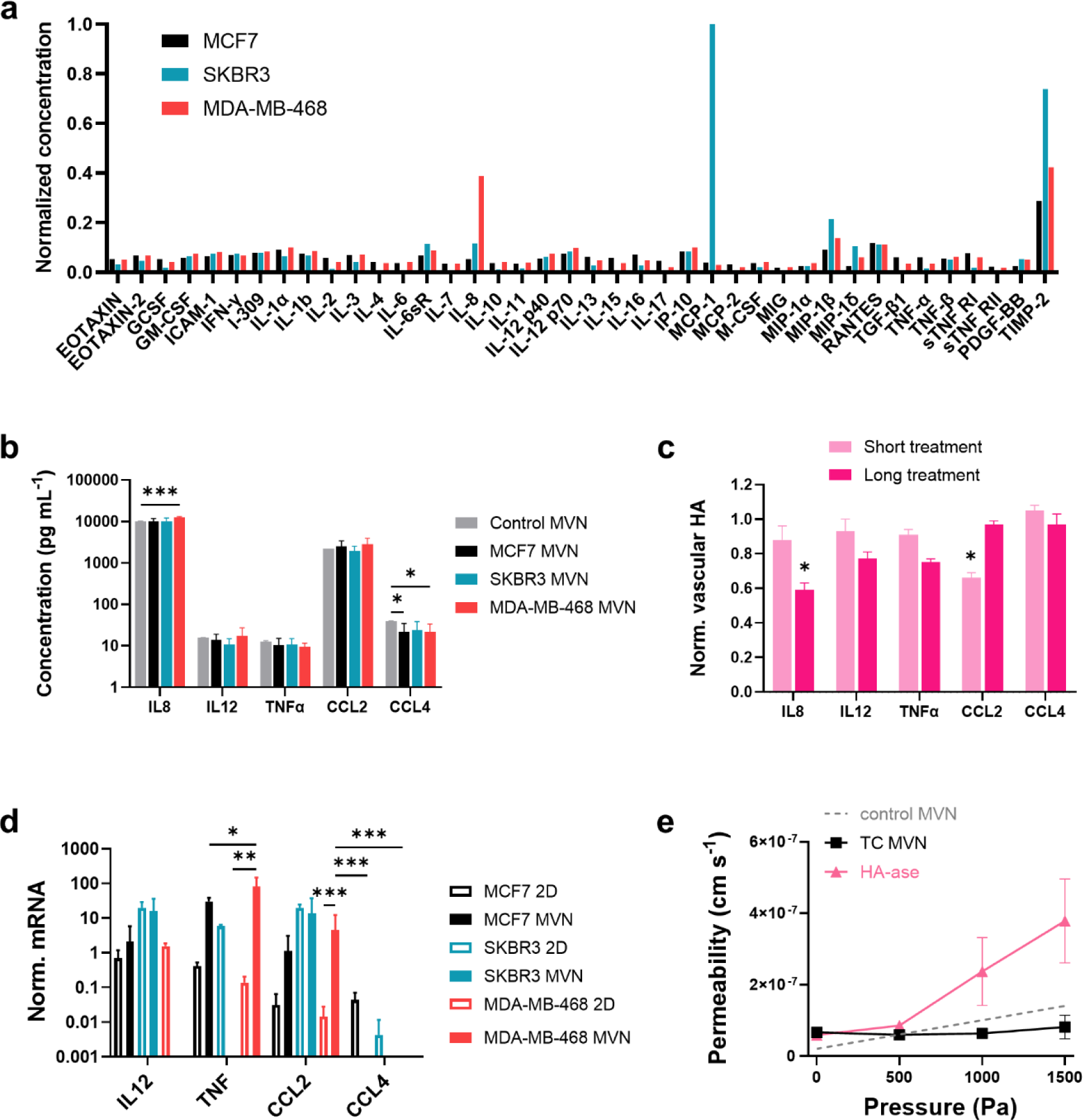
(a) Pro-inflammatory cytokine array in TC-conditioned Vasculife medium. (b) Cytokine concentration in supernatant collected from control and tumoroid MVN devices; n = 3. (c) Quantification of MVN HA by confocal microscopy, normalized to actin signal; n = 3. (d) Gene expression for TCs cultured in 2D or MVNs for genes associated with cytokine production; n =3. (e) Effective permeability of MDA-MB-468 MVNs after treatment with HA-ase; n = 3. Significance assessed by one-way ANOVA; p < 0.5 *, p < 0.01 **, p < 0.001 ***, p < 0.0001 ****.

**Supplementary Figure 8.**
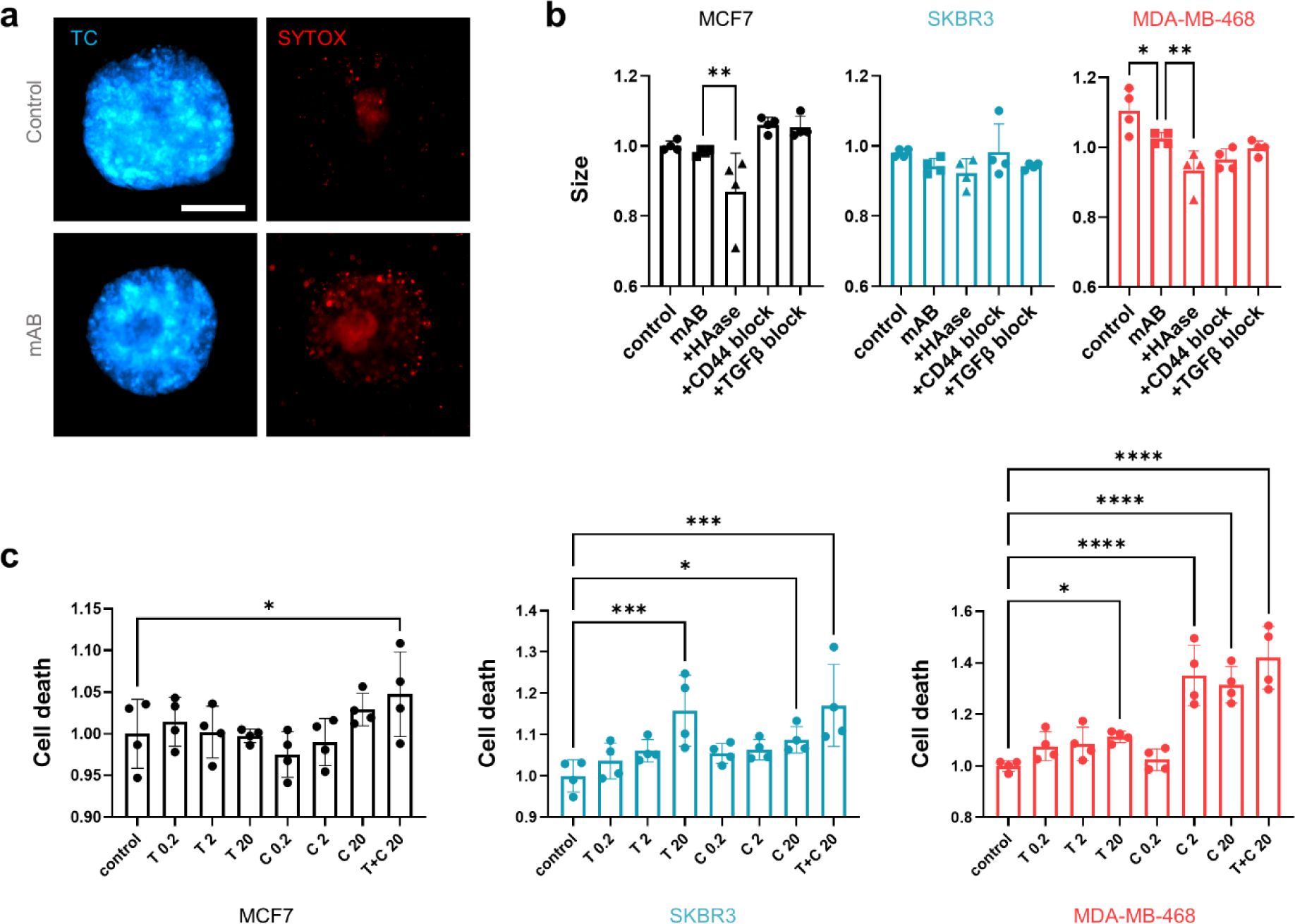
(a) Confocal microscopy image of cell death signal in an MCF7 tumoroid in the MVN device after treatment with Trastuzumab and Cetuximab (“mAB”). (b) Change in size of TC line tumoroids after combination mAB treatment with stromal HA-targeting therapeutic strategies. (c) Normalized cell death signal in TC line tumoroids in well plates treated with Trastuzumab (“T”), Cetuximab (“C”), or both (“T+C”). The concentration indicted is in µg mL^-1^. Significance assessed by one-way ANOVA; p < 0.5 *, p < 0.001 ***, p < 0.0001 ****.

**Supplementary Figure 9.**
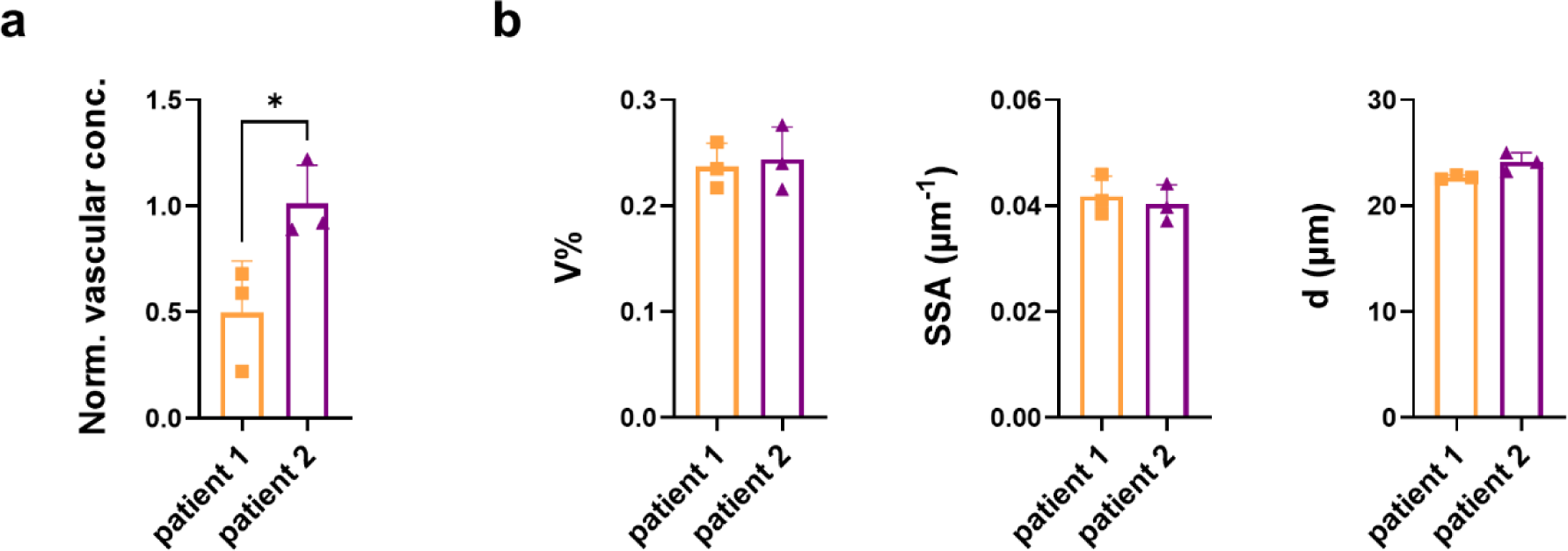
(a) Dextran concentration in primary TC tumoroid MVNs normalized to vessels > 5 mm from the tumoroids. (b) Morphological analysis of primary TC tumoroid MVNs. Significance assessed by student’s t-test; p < 0.5 *.

**Supplementary Table 1.**
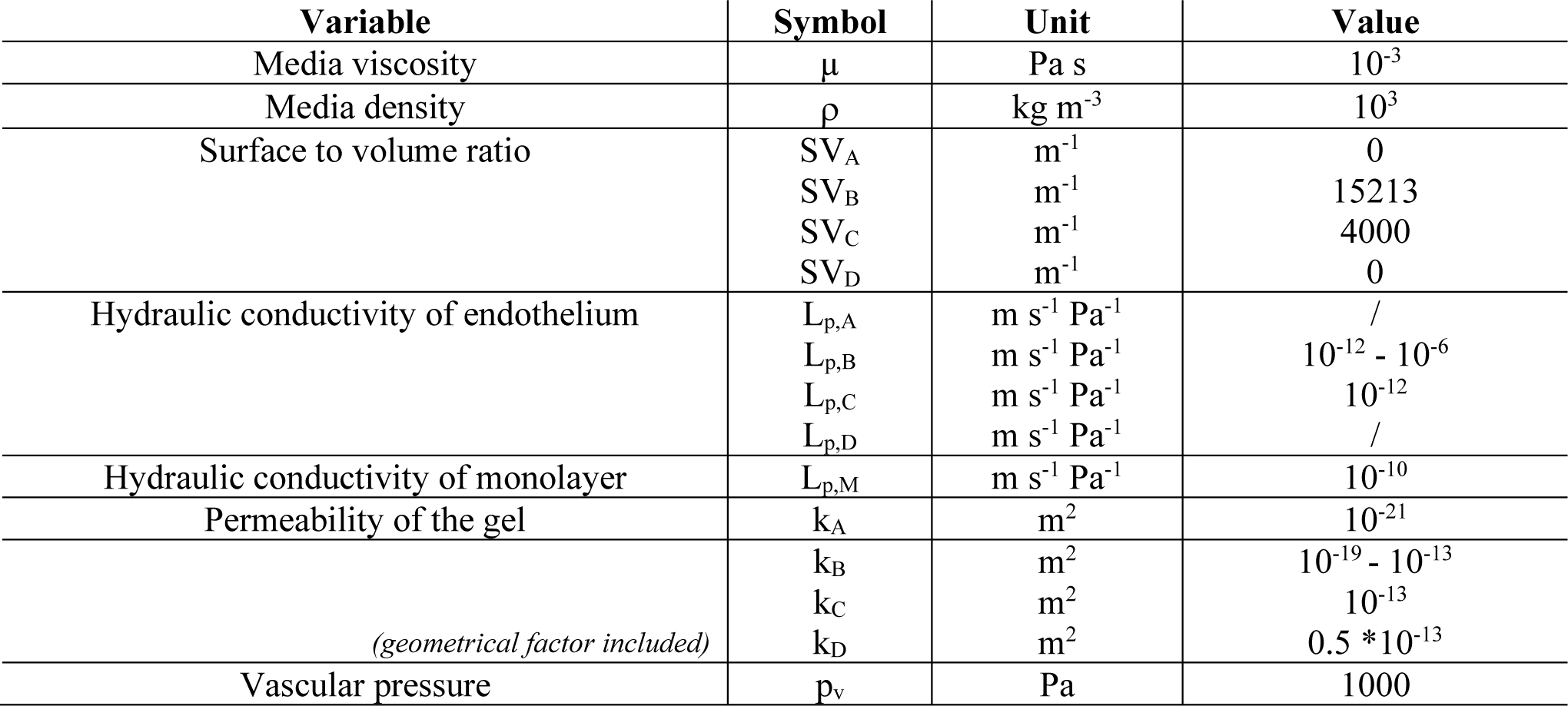
Parameters values used for the computational model analysis.

